# Inhibition of NFAT promotes loss of tissue resident uterine natural killer cells and attendant pregnancy complications in humans

**DOI:** 10.1101/2024.03.07.583906

**Authors:** Rebecca Asiimwe, Brittney Knott, Morgan E. Greene, Emma Wright, Markayla Bell, Daniel Epstein, Stefani D. Yates, Michael V. Gonzalez, Samantha Fry, Emily Boydston, Stephanie Clevenger, Jayme E. Locke, Brian E. Brocato, Constantine M. Burgan, Richard Burney, Nitin Arora, Virginia E. Duncan, Holly E. Richter, Deidre Gunn, Aharon G. Freud, Shawn C. Little, Paige M. Porrett

**Author notes:** Corresponding author: Paige M. Porrett.

## Abstract

Uterine natural killer cells (uNKs) are a tissue resident lymphocyte population that are critical for pregnancy success. Although mouse models have demonstrated that NK deficiency results in abnormal placentation and poor pregnancy outcomes, the generalizability of this knowledge to humans remains unclear. Here we identify uterus transplant (UTx) recipients as a human population with reduced uNK cells and altered pregnancy phenotypes. We show that the NK reduction in UTx correlates with impaired transcriptional programming of NK tissue residency arising from the inhibition of NFAT-mediated signaling. Our observations suggest that NFAT-dependent genes modulate multiple molecular tissue residency programs in uNKs. These include early residency programs involving AP-1-family transcription factors and TGF-β-mediated upregulation of surface integrins. Collectively, our data identify a previously undescribed role for NFAT in uterine NK tissue residency and provide novel mechanistic insights into the biologic basis of pregnancy complications due to alteration of tissue resident NK subsets in humans.

**One Sentence Summary:** Role of NFAT in uterine NK cell tissue residency

## INTRODUCTION

Pre-eclampsia and other diseases of placental malperfusion remain prevalent and impactful problems which complicate approximately 5% of pregnancies in the general population (*1*) Despite decades of intense investigation, our understanding of the mechanistic basis of these diseases is relatively poor, despite a growing appreciation for some of the cellular disturbances which may promote abnormal placentation (*2*). Altogether, these knowledge gaps have severely limited the development of diagnostic testing or therapeutics. Expeditious delivery remains the primary intervention for affected patients, but variability in clinical presentation, tempo, and disease severity often complicates diagnosis (*3, 4*). Consequently, treatment can be delayed, which may lead to severe morbidity or mortality for mother and fetus.

Disordered placentation is thought to contribute significantly to the pathogenesis of pre-eclampsia, and incomplete transformation of the uterine spiral arteries may be foundational in the development of disease. Remodeling of the uterine spiral arteries by invading trophoblast is a key event during placentation, as this process transforms the uterine vasculature into a low pressure, high capacitance system which augments maternal blood flow to the placenta (*5, 6*). In humans, incomplete spiral artery remodeling is associated with structural placental abnormalities and clinical syndromes including pre-eclampsia (*7, 8*). Importantly, mouse models have established that disruptions of uterine natural killer (uNK) cell number or function result in pregnancy complications or loss, often due to abnormal placentation associated with poor spiral artery remodeling (*9–12*). These studies and others thus clearly implicate uNKs in the physiology of uterine spiral artery remodeling and placentation (*5, 9–11, 13–25*). However, differences in the reproductive biology of humans and mice limit generalization of our knowledge from mouse models to human disease (*26*) and pregnancy outcomes have not been well studied in humans with known defects in NK cell number or function. Hence, the specific contribution of uNK cells to pregnancy complications in humans remains unclear.

Given the limitations of mouse models to predict human disease, it is likely that studies of patients at high risk for pre-eclampsia would yield critical insights into disease pathogenesis in humans. Although several risk factors for pre-eclampsia have been described (*27*) the identification of at-risk patients prior to pregnancy can be challenging due to the poor predictive capacity of known risk factors. However, organ transplant recipients represent an immunomodulated population who have reduced numbers of circulating NK cells (*28, 29*) and rates of pre-eclampsia that are six-fold greater than the general population. Moreover, organ transplant recipients have elevated rates of intrauterine growth restriction during pregnancy as well as histologic evidence of placental ischemia (*30–36*). Despite the prevalence of pregnancy complications in organ transplant recipients, the biologic basis for this increased risk is unknown (*37*). Studies of uNKs in this at-risk population are notably absent, likely because of challenges associated with acquisition of samples and the frequency with which pregnancy is discouraged in this population (*38, 39*). Many of these barriers to study may be overcome in the setting of uterus transplantation, where transplantation occurs for the sole purpose of achieving pregnancy and live birth in patients suffering from absolute uterine factor infertility due to congenital absence of the uterus or a history of surgical removal (*40, 41*). Approximately 120 uterus transplants (UTx) have been performed in the world since the first live birth occurred 10 years ago in Gothenburg, Sweden (*42*), with excellent patient and graft outcomes overall (*40, 41, 43–46*). Reproductive and obstetric outcomes have also been excellent, with birth rates comparable to other patient populations utilizing assisted reproductive technologies (*43, 47, 48*). While UTx patients have historically received intense multidisciplinary care at academic medical centers given their overall complexity, excellent clinical outcomes have propelled rapid expansion of the field such that >85% of the transplants performed in the United States since 2020 have occurred at clinical programs and not clinical trials (*49, 50*). Similar to all other solid organ transplant recipients, uterus transplant recipients receive pharmacologic immunosuppression to prevent rejection. Accordingly, the rate of pre-eclampsia in UTx (∼30%) mirrors that of other organ transplant recipients (*40, 41, 51*), suggesting that these individuals represent an opportunity to better understand the immune pathogenesis of pregnancy complications.

In this study, we report the clinical, pathologic, radiologic, and immunologic outcomes of five recipients of a deceased donor uterus transplant, including three with successful live births. Early post-transplant outcomes were excellent, as perfusion of the graft was maintained in all patients, and all patients resumed menstrual cycles within ∼6 weeks of transplant. Complementary single cell and imaging approaches were used to study uNKs derived from endometrial biopsies and term deciduas of UTx; these samples were compared to healthy control (HC) samples that were matched for menstrual cycle phase and gestational age. We found that UTx recipients possessed a reduced frequency of uterine tissue resident natural killer cells (trNKs). Reduced frequency correlated with clinical pregnancy complications and histologic placental abnormalities. Transcriptional programs of tissue residency in UTx trNKs were disrupted compared to HC trNKs. UTx exhibited impaired TGF-β-mediated integrin expression and decreased AP-1-based early tissue residency programs. Through comparative analyses of ex vivo samples and direct pharmacologic inhibition in vitro, we identified a previously undescribed role for NFAT in tissue residency programming of human uNK cells. Importantly, NFAT-dependent residency programs were not equally expressed among uterine trNK subsets, providing a mechanistic explanation for preferential loss of the CD103-expressing trNK3s in UTx recipients. Taken together, these studies link impaired tissue residency programming with defects in uNK abundance to improve our understanding of the pathogenesis of pregnancy complications in humans.

## RESULTS

### Reduction of decidual NKs in UTx recipients associates with placental histopathology and clinical manifestations of placental ischemia

To study the pathogenesis of pregnancy complications in transplant recipients, we measured uterine recovery, function, and pregnancy outcomes after deceased donor transplantation in five UTx recipients (Fig. 1A, fig. S1A, and table S1). UTx received standard pharmacologic immunosuppression as previously described (*40, 41, 51*); this included perioperative T-cell depletion with anti-thymocyte globulin and maintenance immunosuppression with a calcineurin inhibitor (i.e., FK506 / tacrolimus), an anti-proliferative agent (i.e., azathioprine), and an oral steroid (i.e., prednisone). All uterine allografts demonstrated evidence of excellent perfusion after transplantation (Fig. 1, B and C), with resistive indices in the uterine arteries comparable to those in renal arteries of kidney transplants (Fig. 1B) (*52–54*) Although graft perfusion was not assessed routinely at later time points after transplantation, additional imaging in one recipient 55 days after transplantation demonstrated continued excellent graft perfusion (fig. S1B). All recipients began regular menstrual cycles within 6 weeks of transplantation (40.8 ± 21.2 days [mean±SD]; median = 34 days; range = 25-78 days) (Fig. 1D), indicating physiologic recovery and synchronization of the allograft with the recipient’s hypothalamic-pituitary-ovarian hormonal axis. 5 of 9 embryo transfers resulted in pregnancy, with 4 of 9 resulting in a live birth (n=3) or ongoing pregnancy >35 weeks (Fig. 1A). All infants were delivered by cesarean section at 33.6 weeks (UTx3), 34.3 (UTx1) and 36.4 weeks (UTx2) of gestational age due to maternal indication (Fig. 1, A and E). Despite early delivery, infants had good Apgar scores and appropriate weights for gestational age (Fig. 1E). No infants required respiratory support, and length of stay in the neonatal intensive care unit was brief for all neonates (4 days, 0 days, and 3 days, for UTx1, UTx2, and UTx3, respectively).

**Fig. 1.**
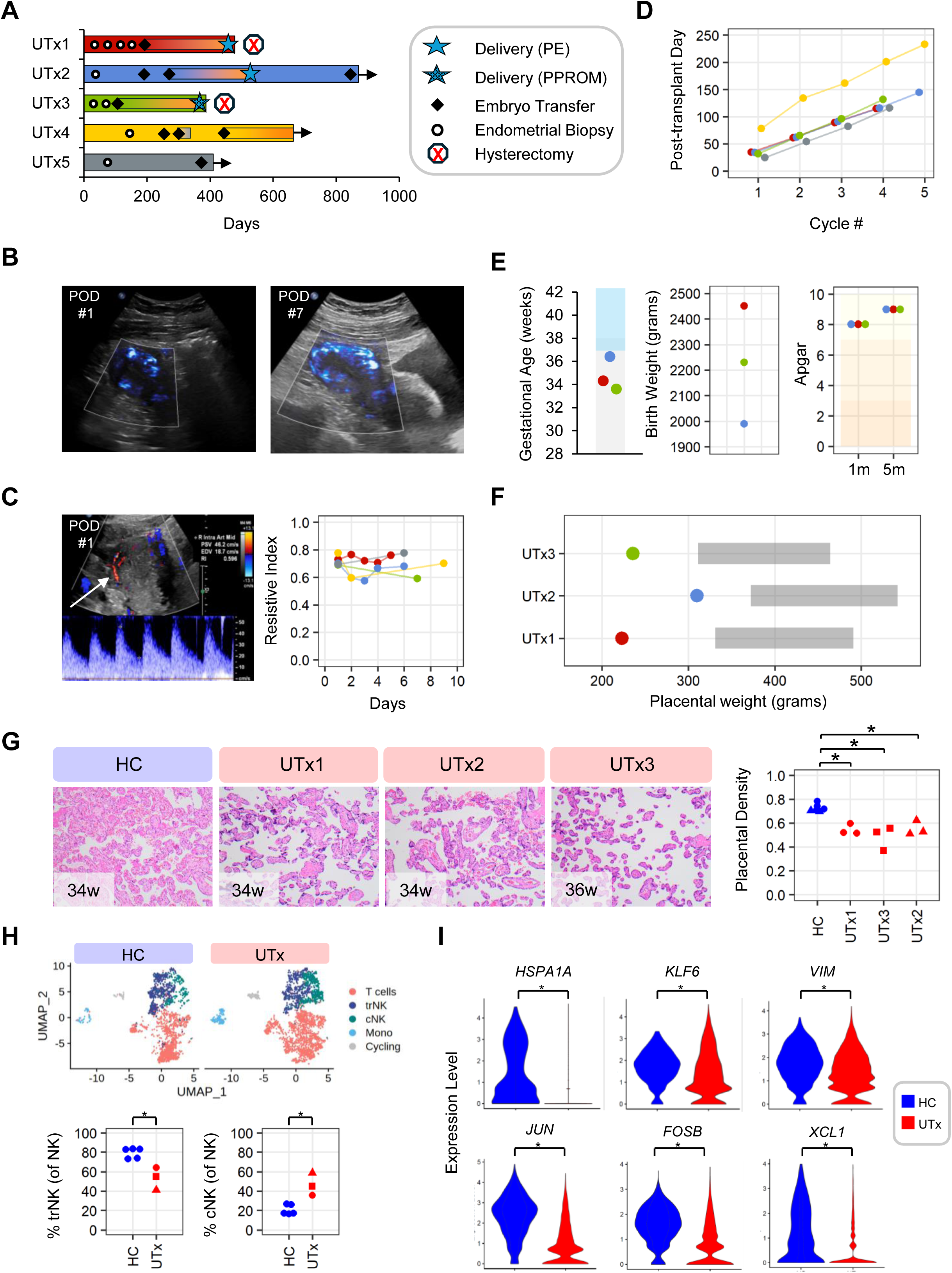
Outcomes of uterus transplant recipients and their pregnancies. (**A**) Graphical summary of clinical course of five recipients of a deceased donor uterus transplant (UTx). Rectangles = active pregnancy. Cesarean deliveries are indicated by stars and were performed prior to 37 weeks for maternal indication. Hysterectomies in UTx1 & UTx3 occurred at time of delivery. PE = pre-eclampsia. PPROM = preterm premature rupture of membranes. ET = embryo transfer. (**B**) Intraparenchymal perfusion of uterine graft (UTx2) on routine surveillance ultrasound within first week after UTx. *<u>Left</u>*: Long axis view of the uterine transplant allograft at the mid-portion, with uterine graft delineated in the white box. MicroFlow Imaging (MFI) Doppler demonstrates scattered vascularity/perfusion throughout the visualized portions of the graft (blue coloring). POD = post-operative day. *<u>Right</u>*: Long axis view of the uterine transplant allograft at the cranial/fundal aspect. MicroFlow Imaging (MFI) Doppler demonstrates robust vascularity/perfusion throughout the visualized portions of the graft (blue coloring) with more extensive vascular networks visible on POD6 than on POD1 (*left*). (**C**) Measurement of intra-arterial flow in a uterine allograft by Doppler ultrasound on POD1 (UTx1). *<u>Left</u>*: Color and spectral Doppler image of the uterine allograft centered on a right sided intraparenchymal uterine artery in the mid-uterus. Color Doppler demonstrates the intrauterine blood vessels (arrow), and spectral Doppler evaluation (bottom, blue wave form) reveals normal low-resistance intrauterine waveforms with expected brisk systolic peaks and appropriate forward diastolic flow. *<u>Right</u>*: Plot of resistive index measured in the right uterine artery at the midbody for each recipient (Mean RI = 0.69 ± 0.07 [SD]). Line colors map to individual recipients as in (A). (**D**) Resumption of menstrual cycles early after transplantation. (**E**) Neonatal outcomes after delivery in three UTx. *<u>Left</u>*: Gestational age at delivery. Gray box indicates range of gestational age of UTx births reported in the literature. Yellow box indicates range of gestational age considered as term pregnancy. UTx1=red; UTx2=blue; UTx3=green. *<u>Middle</u>*: Birth weight in grams. *<u>Right</u>*: Apgar scores at 1 and 5 minutes. Orange shaded rectangles indicate range of Apgar scores and are defined as “reassuring” (7–10), “moderately abnormal” (4–6), and those which indicate need for additional resuscitation (0–3). (**F**) Placental weight in grams for UTx deliveries. Gray boxes indicates expected range for gestational age. (**G)** Placental histology of UTx recipients (H&E 10X). *<u>Left</u>*: Representative sections through the basal plate reveal villi that are sparsely distributed, markedly similar in caliber, and show less branching and have increased numbers of syncytial knots relative to control placenta (HC). Gestational age is indicated. A 36 week placenta from a healthy control was also analyzed (not shown). Sections were taken through the mid basal plate. *<u>Right</u>:* Density of placental villi is plotted for each sample as shown. Density was calculated as the number of pixels covered by placental villi. Individual data points represent technical replicates of two control placentae (34w [blue circle] + 36w [blue triangle]) and 3 UTx placentae. *P-value <0.018 (One-sided Wilcoxon test). UTx1 and UTx3 are compared to 34w HC. UTx2 was compared to 36w HC. (**H**) UMAP visualization of NK and T cells re-clustered from CD45+ cells FACS-sorted from term decidua from three uterus transplant recipients and aggregated with reference immune cell scRNA-seq data from five healthy control term decidua (HC) [HC= 2,285 cells; UTx=2,445 (downsampled)] (see fig. S1). *P-val<0.018 (one-tailed Wilcoxon test) (**I**) Expression of selected genes associated with tissue residency in tissue resident NK cells from decidua of HC and UTx (clusters shown in (H)). *P-val<2E-16 (Wilcoxon test).

Despite the ability to achieve pregnancy and live birth, all three UTx deliveries were associated with clinical phenotypes of placental maternal vascular malperfusion (*55, 56*). Notably, UTx1 and UTx2 developed pre-eclampsia with elevated blood pressure and serum creatinine, while UTx3 developed preterm premature rupture of membranes without evidence of acute chorioamnionitis. All UTx placentas were small for gestational age (Fig 1F), and all placentas had histologic evidence of maternal vascular malperfusion, including patchy regions of increased syncytial knots and slender, sparse villi with widening of the intervillous space consistent with distal villous hypoplasia and accelerated villous maturation (Fig. 1G) (*7, 55–57*). While these placental histopathologic findings were in line with a prior study of placentas from kidney transplant recipients (*36*) we noted no evidence of decidual arteriopathy in UTx placentae (fig. S1C).

Given the lack of histologic evidence of endothelial dysfunction, we searched for other causes of aberrant placentation in UTx. Considering the critical role played by immune cells in placentation, we hypothesized that decidual immune populations might be altered in these immunosuppressed individuals. We thus performed scRNA-seq on CD45+ immune cells sorted from UTx deciduas (n=3) and compared these data with CD45+ cells sorted from healthy controls (n=5) (*58*). We utilized term decidua for this comparison as decidual immune populations are dynamic and change considerably over gestation (*58–60*). Consistent with the work of others (*58*), we found that monocytes and T cells were the most frequent immune cells in term decidua, followed by NK cells and B cells (fig. S1D). Although the distribution of major immune populations was not different between HC and UTx term deciduas (fig. S1D), we identified and re-clustered decidual NK cells for further analysis given the expected low frequency of these cells at term (fig. S1, D and E) (*58*). Among decidual NK cell populations, we were able to identify dNKs expressing genes associated with tissue residency (fig. S1F), along with cells expressing *FCGR3A* (fig. S1E), as previously reported (*58*). In contrast to tissue resident dNKs (trNKs), *FCGR3A*-expressing cells possessed cytotoxic gene programs and were thus identified as conventional NK cells (cNKs) (fig. S1F). Although trNKs were the most abundant group of NKs in control term decidua, cNK frequency was significantly higher in UTx than controls (Fig. 1G). Comparison of genes associated with tissue residency (e.g., *VIM*, *KLF6*, etc.) between HC and UTx trNK suggested a significant depression of tissue residency transcriptional programming in UTx decidual trNK (Fig. 1H). To determine whether specific trNK subsets were affected in UTx deciduas, we identified canonical dNK2 and dNK3 cells among trNKs by expression of reference gene signatures (fig. S1G) (*61*). dNK1 cells were absent in healthy control term decidua (fig. S1G), consistent with prior findings (*58*). Of the remaining populations, we found dNK2 trNKs were preserved between HC and UTx (fig. S1H), whereas dNK3s were significantly reduced in UTx vs HC (fig. S1H), suggesting that the reduction in tissue resident decidual NKs in UTx appeared to most affect the dNK3 subset.

### Reduced frequency of NK cells among endometrial immune cells in UTx

Given that NK cells are recruited into the uterus and develop residency before expanding during pregnancy (*62–64*), we hypothesized that NK cells were diminished prior to pregnancy at earlier time points in UTx after transplantation. We thus performed endometrial biopsies on five UTx >3 months after transplantation who had resumed normal menstrual cycling (9 biopsies; 6 in secretory phase) and compared these results to six HC volunteers (table S1). Given the variable expression of CD56 across NK cell subsets, we assured capture of all NK cells by sorting CD45+ cells from endometrial biopsies and performing scRNA-seq (Fig. 2A and fig. S2A). After data processing and Harmony integration (*65*), we analyzed ∼96,500 CD45+ endometrial immune cells from 15 biopsies (fig. S2, A and B), which included various lymphoid and myeloid populations such as NK cells, T cells, group 3 innate lymphocyte cells (ILC3s), B cells, mast cells, plasmacytoid dendritic cells, and monocytes/macrophages (Fig. 2A and B).

**Fig. 2.**
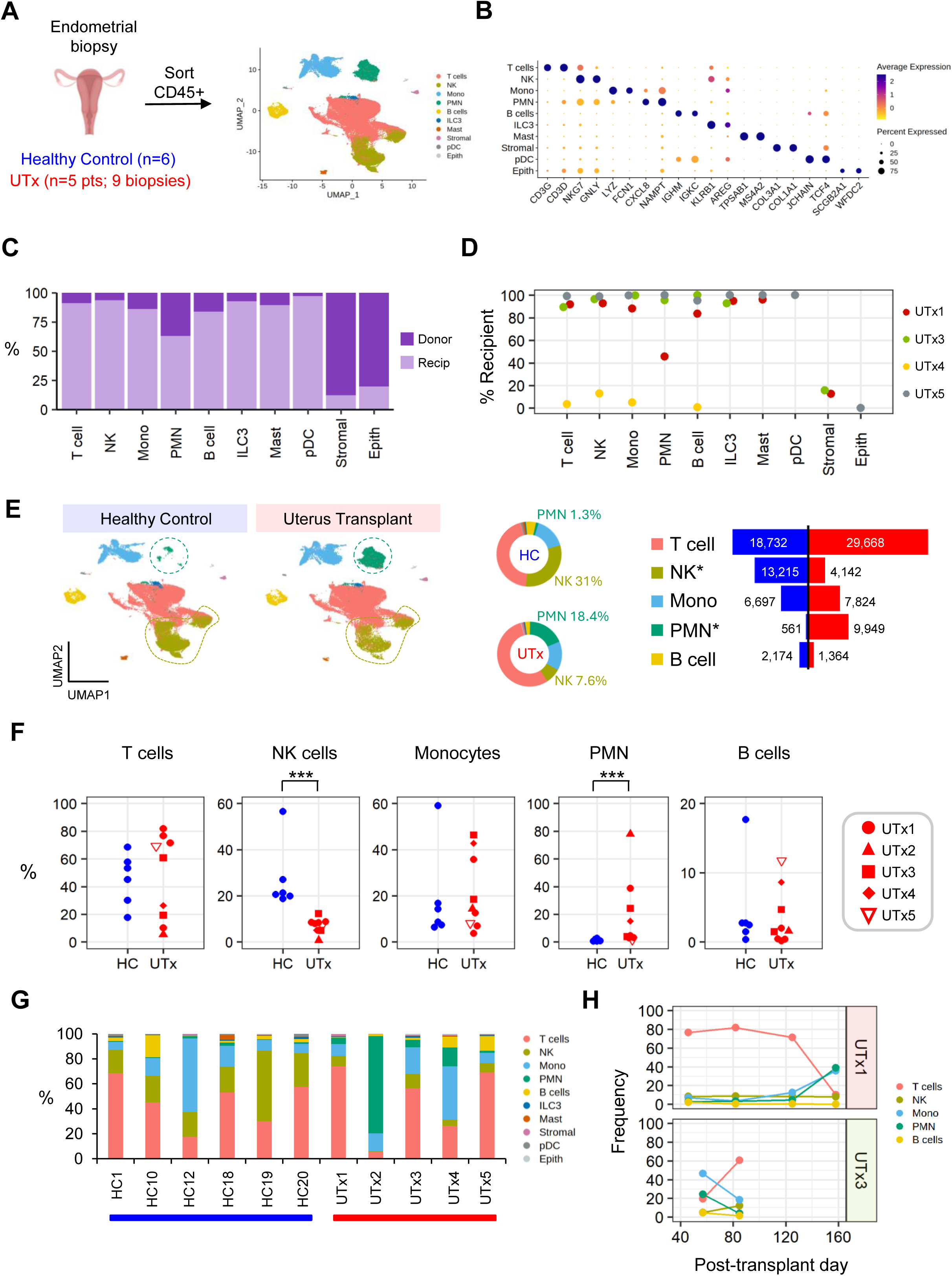
Reduced NK cell frequency among UTx endometrial immune cells. **(A)** UMAP visualization of 96,695 cells FACS-sorted from endometrial biopsies from six healthy control volunteers (HC) (secretory phase) and five uterus transplant recipients (UTx) (6 secretory phase and 3 proliferative phase; 9 total biopsies) (UTx). (**B**) Expression of cell-type specific marker genes among sorted cells. (**C**) Origin of individual cell types in UTx endometria. Genotype of donor and recipient was determined by whole exome sequencing pre-transplant donor and recipient PBMCs for four of five UTx donor:recipient pairs (see Methods). (**D**) Frequency of recipient-derived cell types by individual UTx recipient. Cells were pooled for individuals with multiple biopsies (i.e., UTx1 & UTx3). Genotype is indicated for a given immune population only if a sample had >20 counts of that cell type. (**E**) Distribution of immune cells in HC vs UTx endometrium (**F**) Distribution of immune cells in HC vs UTx endometrium by cell type. ***P<0.001; One-tailed Wilcoxon test. (**G**) Frequency of cell types per individual sample. (**H**) Changes in cell type frequency over time from two UTx with serial biopsies.

In line with prior work (*46*), immune cells in uterus transplants were largely recipient-derived (Fig. 2C and D), confirming that all major immune lineages can be reconstituted in uterus transplant recipients from the peripheral blood. However, immune cells in one recipient (UTx04) were primarily donor-derived (Fig. 2D), a variance which we confirmed by study of endometrial biopsies taken of the donor uterus prior to procurement (Fig. S2B). Notably, endometrial biopsies of the donor organ prior to procurement and transplantation served as critical controls to ensure the accuracy of our genotyping (fig. S2B). Comparison of HC and UTx immune frequencies revealed an altered distribution of immune cells in UTx (Fig. 2E and F), with an increase in polymorphonuclear leukocytes (PMNs; neutrophils) and a decrease in the frequency of NK cells. While PMNs were not increased in all UTx biopsies (Fig. 2F) or all individual UTx recipients (Fig. 2G), we found that NK cells were reduced in frequency in all UTx recipients (Fig. 2F and G), and this reduction was stable across time in longitudinal biopsies despite variation in other immune cell frequencies (Fig. 2H). To validate these results, we used flow cytometry to compare the frequency of uNK cells between UTx and HCs. Although tissue for flow cytometric analysis was available from only three of the nine UTx biopsies, these analyses reproduced the findings of our single-cell analyses, as we observed a statistically significant reduction in NK cell frequency among endometrial immune cells in UTx versus HCs, independent of menstrual cycle phase (fig. S2C).

### Reduced NK cell density in UTx endometrium

The reduction in NK cell frequency among CD45+ endometrial immune cells in UTx could result from either a reduction in NK cell density in the tissue or from the presence of excess CD45+ non-NK cells. To determine whether NK cell density was reduced in UTx endometrium, we performed immunofluorescence microscopy to evaluate multiple fields of sections of endometrium from UTx and HCs. Biopsy sections (Fig. 3A) were stained with DAPI and anti-CD56 and data collected by epifluorescence (Fig. 3, B to F) at high resolution. Nuclei were segmented by custom semi-automated processing of the DAPI channel (see Methods) and assigned a status of positive or negative for CD56 expression (Fig. 3C and D). Multiple independent image fields were collected, each representing an area of about 0.13 mm^2^ and containing 870±128 (mean ± SD) cells per field. The fraction of CD56+ cells was then calculated for each field (Fig. 3G to I).

**Fig. 3.**
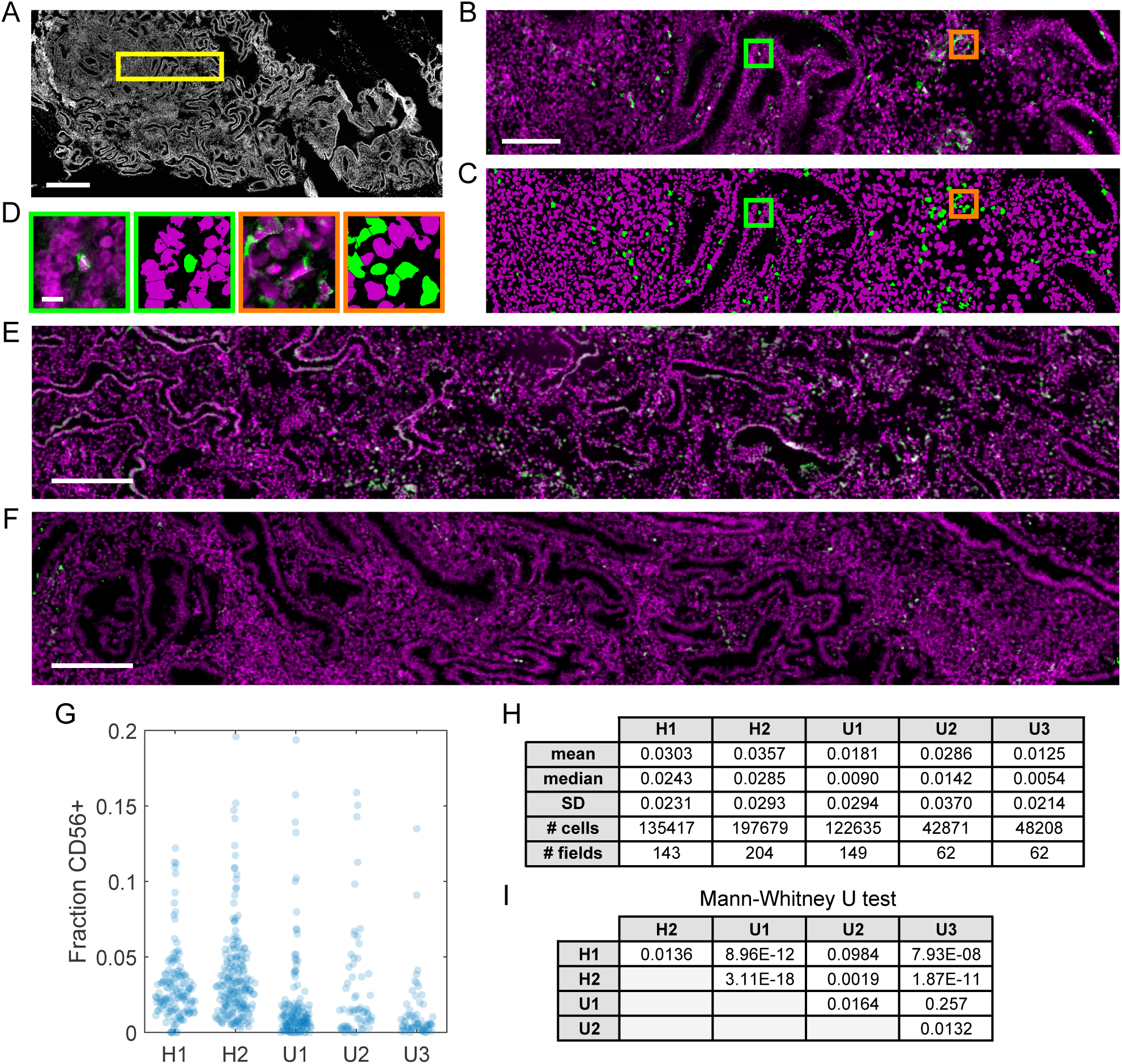
CD56+ cells are reduced in endometrial tissue sections of transplanted uteri. (**A**) Low-magnification DAPI channel from fluorescence microscopy of healthy control endometrial biopsy (“H1” in panels G-I). Box (yellow) indicates area shown in panel B. (**B**) High magnification image showing DAPI (magenta) and anti-CD56 staining (green). (**C**) Computerized rendering of panel B showing CD45+ (green) and CD56- (magenta) nuclei. Boxes indicate areas shown in D. (**D**) Data (left) and computer rendering (right) of the green and orange boxed areas shown in B and C. (**E** and **F**) DAPI (magenta) and anti-CD56 (green) staining in a second control biopsy (E) and a biopsy taken 57 days after uterus transplant (F), designated “H2” and “U1”, respectively, in panels G-I. (**G**) Fraction of nuclei designated CD56+ in two healthy control (H1 and H2) and three uterus transplant biopsies (U1-U3). Data points represent individual 0.15 mm2 imaging fields. (**H**) Summary statistics for the five biopsies. (**I**) p-values (Mann-Whitney U test) for all pairwise comparisons of the five biopsies. Scale bars 500 µm (A), 100 µm (B), 10 µm (D), 200 µm (E,F). See table S1 for additional details regarding samples.

CD56+ cells were observed either as isolated individuals or in groups (Fig. 3D). Counts of cells per field varied widely within the same biopsy (Fig. 3G), suggesting non-homogenous distribution of spatial cues directing NK localization in the tissue in both HC and UTx endometrium. By comparing the median values of the fraction CD56+ cells, we found that NK cell density was reduced by roughly a factor of three in UTx biopsies (2.6% in HC versus 0.9% in UTx). p-values resulting from pairwise Mann-Whitney U tests showed that UTx samples tended to differ more from control than from each other (Fig. 3I). In contrast, the same analysis using anti-CD3 staining reveals that the density of CD3+ cells is reduced by about 40% in UTx compared to control (11% in control versus 7% in UTx; fig. S3). We conclude that NK cell density is reduced in UTx, supporting the observations from flow cytometry and scRNA-seq analyses.

### Reduction of tissue resident NK subsets in uterus transplant endometrium

To investigate the mechanism of NK loss in UTx recipients, we first determined whether the NK cell reduction in UTx endometrium affected all uterine NK subsets equally. Consistent with prior studies (*64, 66, 67*), we identified two major populations of NK cells among endometrial immune cells (Fig. 2A and 4A) which could be distinguished by differential expression of *NCAM1* and *FCGR3A* (fig. S4A) as well as transcriptional programs governing tissue residency and cytotoxic effector function (fig. S4, B to D). Tissue resident NK cells (trNK) expressed integrins (i.e., *ITGA1, ITGAX, ITGAD*) (Fig. 4B and fig. S4B) and chemokine effector molecules (i.e., *XCL1*, *XCL2*) (fig. S4, B and D) previously described for tissue resident lymphocytes (*68–70*). In contrast, conventional NK cells (cNK) expressed a variety of molecules associated with circulating populations, including *S1PR5* and *KLF2* (Fig. 4B and fig. S4, B and C) (*64, 68–73*). Moreover, the endometrial tissue resident cluster expressed a gene signature associated with tissue residency in human lung NK cells (Fig. 4C and table S2) (*71*). Whereas cNK number and frequency were similar in UTx vs. HCs, trNK number and frequency were significantly reduced in UTx (Fig. 4, D and E).

**Figure 4.**
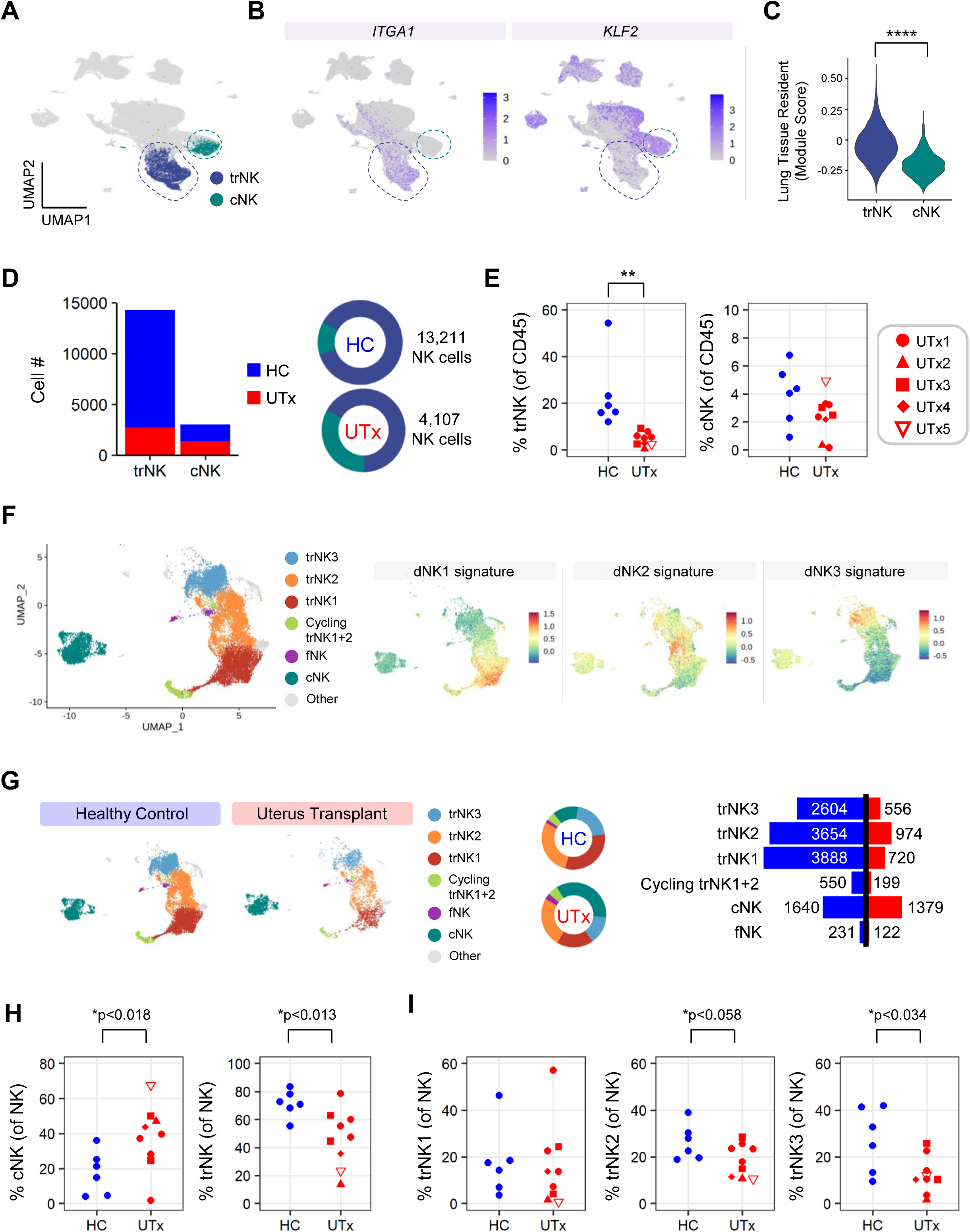
Reduction of trNK subsets in UTx endometrial biopsies. (**A**) Feature plot of tissue resident NK (trNK) and conventional NK (cNK) subsets superimposed on CD45+ UMAP of aggregated immune cells from HC and UTx endometrial biopsies (see Fig. 2). Non-NK populations are shown in gray. (**B**) Reciprocal expression of marker genes of tissue residency (*ITGA1*) and circulation (*KLF2*) in trNK and cNK cells. Dashed lines indicate populations of trNK and cNK. (**C**) Expression of human lung tissue resident module score by endometrial trNKs vs. cNKs. ****P-val < 2E-16. (**D**) *<u>Left</u>*: Total number of trNK and cNK cells in sequenced endometrial biopsies from HC (n=6) and UTx (n=5 pts; 9 biopsies). *<u>Right</u>*: Frequency of NK cells that are trNK and cNK in each group. (**E**) Frequency of trNK and cNK of sequenced CD45+ cells in each biopsy. ***P-val<0.0002 (one-tailed, Wilcoxon). (**F**) *<u>Left</u>*: UMAP projection of 17,318 NK cells re-clustered from trNK and cNK subsets depicted in (A). *<u>Right</u>*: Expression of reference decidual NK signatures (Vento-Tormo, Nature 2018) by aggregated HC + UTx endometrial NK cells. (**G**) *<u>Left</u>*: Split UMAP showing cell density of trNKs in HC and UTx. *<u>Middle</u>*: Frequencies of trNK subsets by group. *<u>Right:</u>* Numbers of cells sequenced in each trNK subset from HC or UTx. (**H**) Frequency of trNK and cNK cells as a fraction of total NK cells in HC and UTx (by biopsy). (**I**) Frequency of major trNK subsets in individual biopsies of HC and UTx.

Because uterine trNKs are transcriptionally and functionally heterogeneous (*61, 64, 74*), we next determined if all trNK subsets were reduced in UTx or whether the trNK defect was limited to specific trNK subsets. To determine the frequency of specific trNK subsets among endometrial immune cells, we re-clustered all endometrial NK cells (Fig. 4F and fig. S4E) and were able to identify major endometrial trNK subsets (trNK1, trNK2, trNK3) among *ITGA1*-expressing trNKs (fig. S4E). Endometrial trNK subsets were identified based on expression of reference decidual NK signatures (dNK1, dNK2, dNK3) (Fig. 4F) (*61*) and marker genes (fig. S4F and table S2). We found that the ratio of trNK to cNK was reversed in UTx compared to HC given the apparent preservation of cNKs in UTx recipients (Fig.4, G and H). While HC and UTx had similar frequencies of trNK1 cells, we found statistically significant reductions in trNK3 frequency in UTx recipients, with a trend towards reduction of trNK2 cells as well (Fig. 4I). Taken altogether, these data suggest that the NK reduction in UTx endometrium is driven primarily by a loss of tissue resident NK cells. These data also suggest that the loss of decidual trNK cells observed in UTx is not unique to pregnancy but instead occurs as a consequence of alteration of uterine tissue resident NK biology.

### Altered transcriptional programming of canonical TGF-β-mediated tissue residency in UTx trNK

Having established that trNK cells were reduced in UTx endometrium compared to HCs, we aimed to understand the molecular mechanisms responsible for this reduction. To test the hypothesis that transcriptional programs of trNK development were altered in UTx endometrial trNK, we performed CITE-seq analysis on CD45+ cells aggregated from three HC and five UTx endometrial biopsies from three individual recipients (table S1 and fig. S5A). CITE-seq enabled simultaneous analysis of surface phenotype and transcriptomes, thus allowing study of changes in both protein and gene expression across populations. NK cells were identified among CD45+ endometrial immune cells based on their transcriptomes as well as surface phenotype (i.e., CD56+CD14-CD3-) (fig. S5) and were re-clustered for downstream analyses (Fig. 5A and fig. S6A). As in our prior analyses, NK cells resolved into two major groups of cNK and trNK subsets (Fig. 5, A and B, and fig. S6, A to D) which were distinguished by differential expression of genes associated with cytotoxicity (fig. S6, B to D) as well as reciprocal expression of CD49a (*ITGA1*) and CD16 (*FCGR3A*) (Fig. 5B, and fig. S6, B and C). cNK and trNK subsets were further distinguished by variable expression of CD56, CD45RA, and KLRG1 among other genes and proteins (fig. S6, B to D). As predicted by our prior analysis based exclusively on the transcriptome (Fig. 4), CD49a+ trNK cells contained cells with high expression of decidual NK signatures and dNK marker genes (fig. S6, E and F) (*61*). We were further able to confirm the surface phenotype of these trNK subsets which was predicted from their transcriptional signatures, with trNK1 cells having the highest expression of KIRS and CD39, while trNK3 had the highest expression of CD103 (Fig. 5B) and CD69 (fig. S6G). Altogether, these analyses facilitated the annotation of these endometrial NK cells as trNK subsets corresponding to those in the decidua (i.e., dNK1 ≈ trNK1, dNK2 ≈ trNK2, and dNK3 ≈ trNK3).

**Figure 5.**
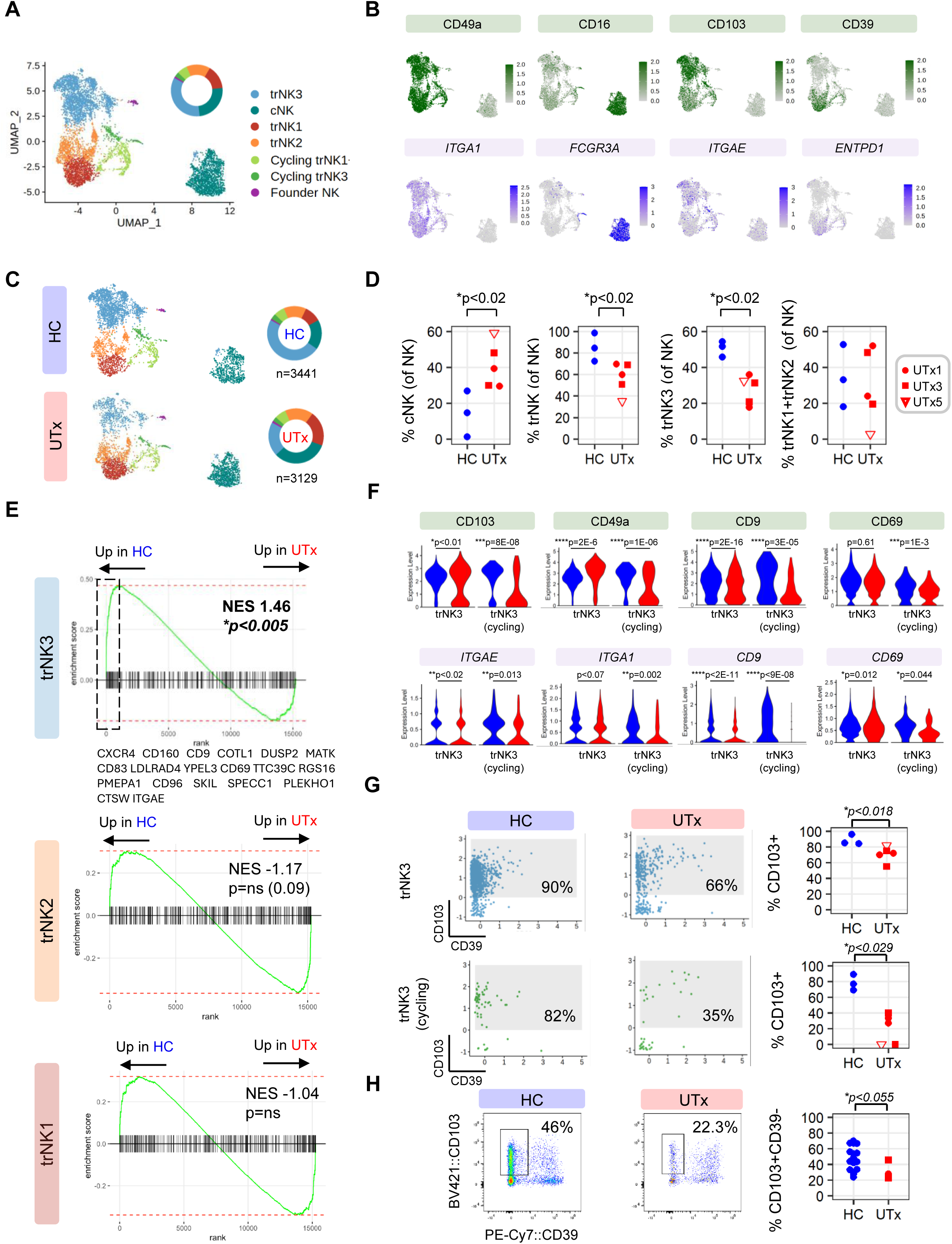
Preferential loss of TGF-β-dependent trNK subsets in UTx endometrium. (**A**) UMAP visualization of 6,570 NK cells re-clustered from CD45 immune cells sorted from 3 HC secretory phase endometrial biopsies and 5 UTx secretory phase endometrial biopsies (3 individual patients), after removal of NKT cells. (**B**) Feature plots of genes and proteins delineating trNK subsets and cNKs in reclustered NK cells. (**C**) Split UMAP of HC and UTx endometrial NK cells indicating differences in frequency of NK subsets between HC & UTx. (**D**) Quantification of differences between trNK frequencies among UTx and HC. (**E**) Gene set enrichment analysis of TGF-β target genes showing enrichment of TGF-β targets in genes upregulated in HC vs. UTx in the trNK3 subset, but not among trNK2 cells or trNK1 cells. (**F**) Comparison of indicated gene and protein expression of selected TGF-β targets among trNK3 and cycling trNK3 subsets in HC and UTx biopsies. (**G**) Scatter plot visualization of CITE-seq antibodies against indicated proteins in trNK3 and cycling trNK3 cells from HC and UTx endometrial biopsies. Shaded box indicates CD103+ cells, quantified toward the right. (**H**) Flow cytometric validation of reduced CD103+ trNK3 frequency in UTx recipients. Single cell suspensions of secretory phase biopsies of 16 healthy control volunteers and 2 individual UTx recipients (3 biopsies) were stained and analyzed. *<u>Left</u>*: Representative flow plots are shown. Gated cells represent the frequency of CD103+ CD39-trNK3 cells of parent gated trNKs (live CD45+CD56+CD16-CD49a+ singlets). UTx (n=3; UTx1 [circle], UTx3 [square]). *one-sided Wilcoxon test.

Next, we compared endometrial trNK subsets among HC and UTx samples. As in our prior analyses, we noted that tissue resident subsets were less frequent among UTx CD56+ NK cells compared to healthy controls (Fig. 5, C and D). In line with our analysis of decidual NK cells in UTx recipients, CD103+ trNK3 cells were preferentially and markedly reduced in UTx (Fig. 5D). Given the critical role of TGF-β in CD103+ tissue resident lymphocyte development (*69, 70, 75–77*), we hypothesized that TGF-β target genes were downregulated in trNK3 cells from UTx compared to HCs. Indeed, the TGF-β gene signature was downregulated in UTx trNK3 subsets (Fig. 5E), including CD69, CD9, and the integrins *ITGA1* and *ITGAE* (Fig. 5F). Accordingly, the surface expression of the corresponding proteins was typically also reduced in UTx trNK3 and cycling trNK3 subsets (Fig. 5F), although reductions of gene expression and protein expression did not always have the same magnitude. Altogether, these results suggested that cells which possessed trNK3 transcriptomes in UTx had an overall shift in their surface phenotype toward CD103-CD39-trNK2 subsets, which we confirmed by flow cytometric analysis (Fig. 5H). Collectively, these data suggested that disruption of TGF-β-mediated tissue residency programming in UTx samples resulted in an overall reduction of trNK3 subsets, with attenuation of TGF-β target genes and surface proteins typically associated with tissue residency in the remaining cells.

### Reduced expression of a gene signature associated with early residency in UTx

Although the contribution of TGF-β to tissue residency programming is well established (*69, 70, 75–77*), less is known about the temporal orchestration of transcriptional programs of residency as lymphoid cells seed tissues from the blood. Importantly, recent work in a mouse model of LCMV infection has suggested that an array of immediate early response genes (i.e., *JUN*, *NR4A2*, etc.) are not only expressed early after tissue entry but are required for tissue residence (*78*). Moreover, prior work investigating human uterine NK development has suggested that a similar transcriptional program of early tissue adaptation is upregulated in early endometrial immigrants after arrival from the blood (*64*). We thus tested the hypothesis that trNKs were reduced in UTx due to defective expression of this early residency program (ERP) (n=80 genes) in endometrial trNK cells (table S2).

First, we examined the expression of the tissue resident ERP gene signature in healthy control trNK and cNK subsets. While we expectedly found higher expression of the ERP in healthy control trNKs vs. cNKs (Fig. 6A), we found reciprocal expression of groups of ERP genes within the different trNK subsets (Fig. 6, B and C). More specifically, we found that trNK3s possessed higher expression of *JUN, JUND*, and *JUNB*, while trNK1s had higher expression of *RUNX3*, *BHLHE40*, *FOS*, and *NR4A2* (Fig. 6B). To further analyze utilization of these gene signatures, we classified 69 of 81 ERP genes into one of two ERP gene signatures, with the ERP1 signature defined by higher expression in trNK3 cells and the divergent ERP2 signature defined by higher expression in trNK1 cells (Fig. 6, B and C, and table S2). Interestingly, trNK2s had intermediate expression of both ERP1 and ERP2 signatures between trNK1 and trNK3 cells (Fig. 6, B and C).

**Figure 6.**
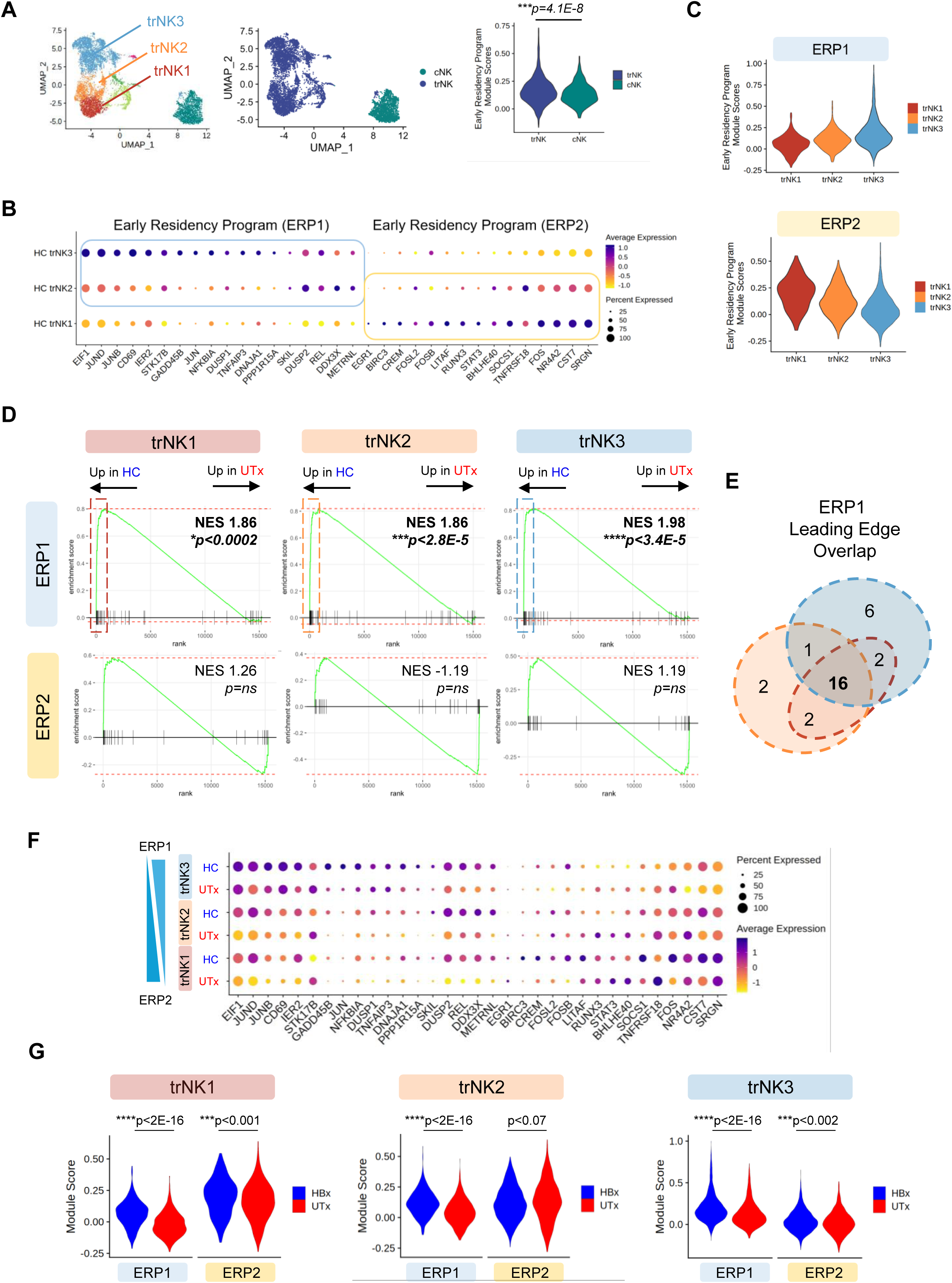
Loss of transcriptional programs of early residency in UTx trNKs. (**A**) *<u>Left & Middle</u>*: UMAP visualization of 6,570 NK cells from 3 HC and 5 UTx endometrial biopsies reproduced from Fig. 5A and fig. S6A for reference. *<u>Right</u>*: Module score of early residency program (n=80 genes) in healthy control trNKs and cNKs (see table S2). (**B**) Expression of top genes of the early residency signature in healthy control trNK subsets. Genes belonging to early residency program 1 (ERP1) and early residency program 2 (ERP2) are indicated. (**C**) Expression of module score of ERP1 (n=41 genes) and ERP2 (n=28 genes) among healthy control endometrial trNK subsets. (**D**) Enrichment of ERP1 and ERP2 gene signatures among genes upregulated in HC vs UTx trNK subsets. Dotted lines represent rank position of leading edge genes (see (**E**)). NES = normalized enrichment score. (**E**) Overlap of ERP1 leading edge genes enriched in trNKs in healthy control samples. See fig. S7 for leading edge gene lists. (**F**) Bubble plot of expression of selected ERP1 and ERP2 genes (as in (**B**)) showing expression in both HC and UTx trNK subsets. (**G**) Violin plots of ERP1 and ERP2 gene signatures in HC vs. UTx (by trNK subset).

Next, we tested the hypothesis that early residency programs were diminished in UTx endometrial NK cells. We performed gene set enrichment analysis of ERP1 and ERP2 signatures in genes differentially expressed in HC vs. UTx trNK subsets. Whereas the ERP2 gene program was not enriched in genes either upregulated or downregulated in healthy control trNKs, we found statistically significant enrichment of the ERP1 gene program in genes upregulated in HCs vs. UTx in all three major endometrial trNK subsets (Fig. 6D). Leading edge genes included members of the JUN AP-1 transcription factor family as well as genes in the NFKB family (fig. S7). Notably, 16 of 41 ERP1 leading edge genes (39%) were enriched in all three trNK subsets (Fig. 6E and fig. S7). Comparison of the most highly expressed individual genes in the two ERP signatures in HC and UTx trNK subsets showcased the magnitude of ERP1 downregulation across trNK subsets (Fig. 6F). We found similar results when comparing module scores calculated for the entire ERP1 and ERP2 gene signatures in HC vs. UTx trNK subsets (Fig. 6G). Collectively, these results suggested that loss of the trNK3 subset in UTx recipients was due in part to the downregulation of a set of early residency genes that was more highly expressed in trNK3s versus other trNK subsets. Although the ERP1 gene signature was downregulated in trNK1s in UTx compared to HC, the early residency gene module which predominated in trNK1 cells (i.e., ERP2) had only modest reductions in expression in UTx, thereby providing a potential explanation for the relative preservation for trNK1 cells in UTx recipients.

### Role for NFAT in tissue residency programming in uterine NK cells

We next considered the mechanism by which transcriptional programming of uterine natural killer cell tissue residency was disrupted in UTx. Because NFAT regulates CD103 gene expression (*79*) and NFAT-dependent genes have been identified within prior studies of tissue resident lymphocytes (*64, 80*), we hypothesized that tissue residency programming was reduced in UTx due to exposure of these individuals to immunosuppression drugs which prevent NFAT nuclear translocation (i.e., calcineurin inhibitors; tacrolimus). We tested for overlap of a curated gene set of NFAT targets (table S2) with genes upregulated during mucosal CD8 T_RM_ differentiation (*78*) and found 42 genes governing lymphocyte tissue residency that could be influenced by NFAT (Fig. 7A). Having identified NFAT-dependent genes involved in the transcriptional programming of tissue residency (i.e., NFAT^TR^ signature), we then tested for enrichment of the NFAT^TR^ signature in endometrial trNK subsets versus conventional NK cells from healthy control volunteers. In line with our prior observations that trNK subsets utilize different tissue residency programs (Fig. 6) (*64*), we found that trNK3 subsets enriched for the NFAT^TR^ signature whereas trNK2 and trNK1 subsets did not (Fig. 7B), given the higher expression of genes such as *CD69* and *JUN* family members in trNK3 cells (Fig. 7C). We next determined whether the NFAT^TR^ signature was differentially expressed in endometrial NK cells from uterus transplant recipients versus healthy controls. We tested for enrichment of the NFAT^TR^ signature among the genes upregulated in HC vs. UTx for each trNK subset. These analyses revealed enrichment of the NFAT^TR^ signature among genes downregulated in UTx compared to HCs in the trNK3 subset (Fig. 7D). Notably, the NFAT^TR^ genes downregulated in UTx trNK3 cells corresponded to genes shared in the early residency program most expressed in trNK3 subsets (i.e., ERP1) (Fig. 7, E and F). Collectively, these data suggest a role for NFAT in tissue residency programming of endometrial trNK cells and identify the largest reduction in the trNK3 subset with highest expression of transcriptional programs that are enriched for NFAT-dependent genes.

**Figure 7:**
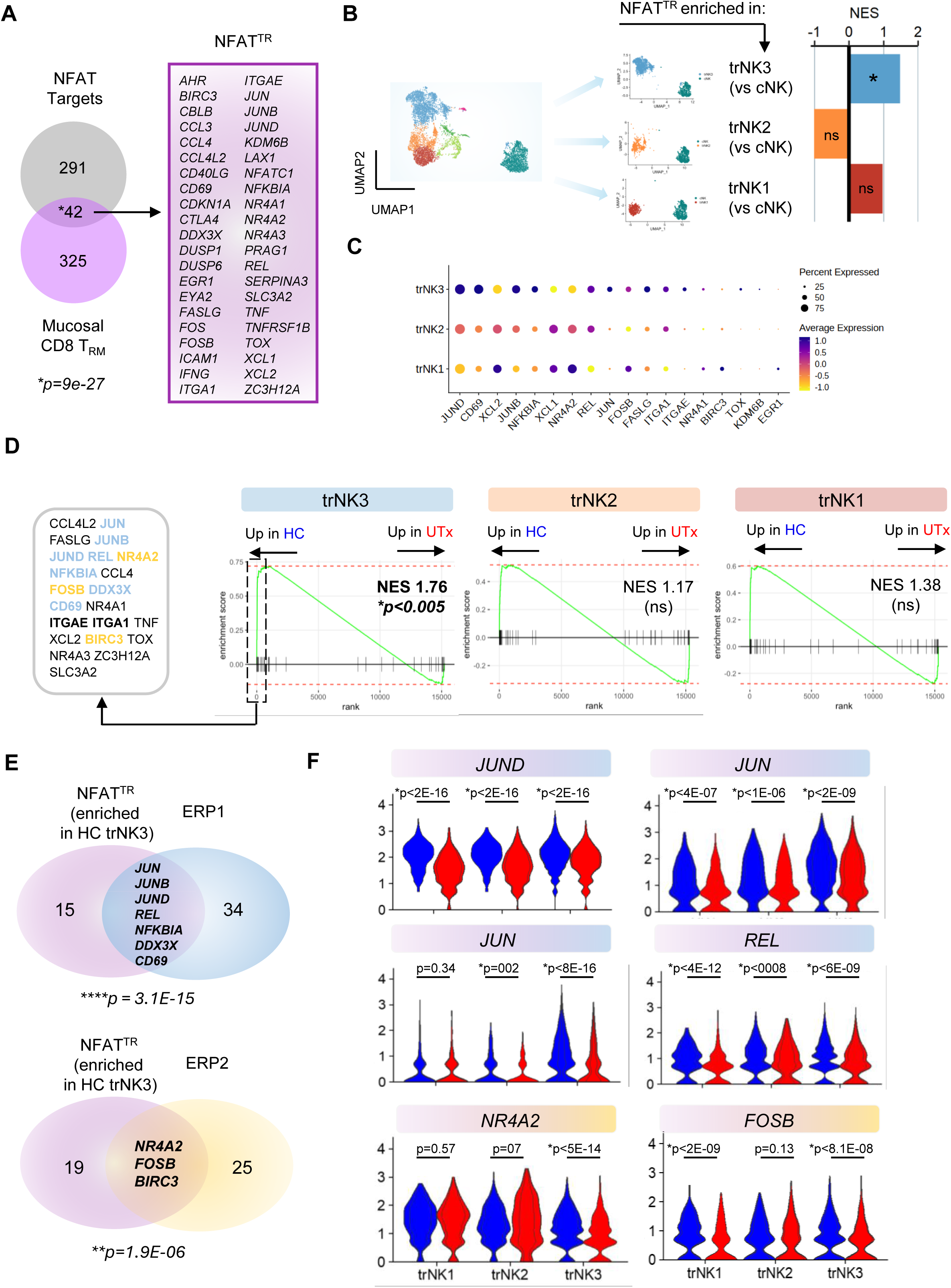
Reduction of NFAT-dependent tissue residency programs in UTx. (**A**) Identification of 42 NFAT-dependent genes involved in tissue residency programming (NFAT^TR^ signature). (**B**) Enrichment of the NFAT^TR^ signature in endometrial trNKs versus cNKs. (**C**) Variable expression of leading edge genes of the NFAT^TR^ across trNK subsets. (**D**) Enrichment of the endometrial NFAT^TR^ signature in healthy control trNK3 subsets. (**E**) Significant overlap of the endometrial NFAT^TR^ signature with the early residency program 1 (ERP1). (**F**) Expression of genes in HC and UTx which overlapped between the NFAT^TR^ signature and early residency programs. Statistical testing: Wilcoxon.

### IL-15 promotes integrin expression in endometrial trNK cells through NFAT

Although NFAT has well-established roles conveying signals downstream of antigen receptors in B and T lymphocytes (*81*), less is known about how innate cells such as NK cells use NFAT to translate environmental inputs. Nevertheless, a small number of prior studies have identified NFAT effects downstream of cytokine receptor signals such as IL-6 (*82*) and IL-15 signals (*83*). Since IL-15 synergizes with TGF-β in NK cells to promote upregulation of integrins important in tissue residency (*84*), we hypothesized that NFAT was contributing to tissue residency programming of trNK cells downstream of IL-15 signals. To test this hypothesis, we tested whether NFAT blockade by FK506 would result in impairment of key integrins on endometrial NK cells in response to IL-15 stimulation. IL-15 stimulation in vitro for 4-7 days was sufficient to induce significant upregulation of CD103 and CD49a expression on endometrial NK cells without the addition of exogenous TGF-β (Fig. 8, A and B). However, the addition of FK506 to these cultures significantly diminished the frequency of CD103+CD49a+ endometrial trNK cells, with concomitant increases in the frequencies of CD103-CD49a+ and CD103-CD49a-cells (Fig. 8, C and D). Altogether, these in vitro studies suggest that cytokine signals which mediate tissue residency programs in endometrial trNK utilize NFAT.

**Figure 8:**
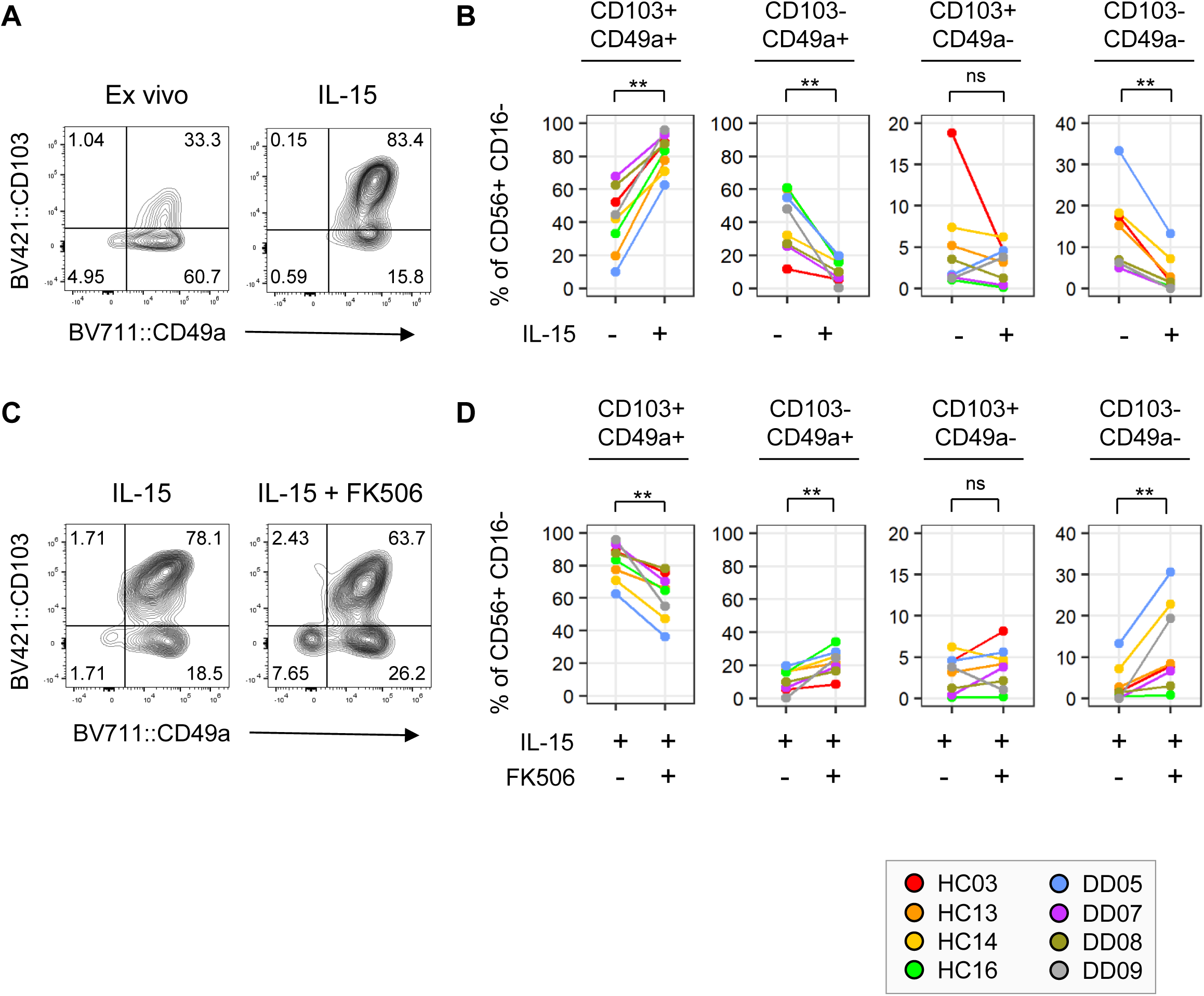
NFAT blockade correlates with impaired IL-15-induced integrin expression on NK cells. Endometrial biopsies from healthy control volunteers (n=4; secretory phase) (HC) and uteri from deceased donors (DD) (n=4; unknown menstrual cycle phase) were collected and single cell suspensions were prepared. For samples analyzed directly ex vivo (**A&B**), an aliquot was stained with fluor-conjugated antibodies and analyzed with a spectral flow cytometer. The remaining aliquots from the same samples were cultured in vitro with 10ng/mL IL-15 +/- 10ng/ML FK506 for 3-6 days in duplicate or triplicate (**C&D**), and analysis was performed with spectral flow cytometry. (**A&B)** Increase in CD49a+ CD103+ trNK3 frequency after IL-15 culture in vitro. **C&D**) Reduction in CD49a+ CD103+ trNK3 frequency after co-culture with FK506 in vitro. Flow panels indicate representative data. **P<0.01; ns=not significant; paired Wilcoxon signed rank exact test.

## DISCUSSION

The primary objective of this work was to improve our understanding of the immunologic basis of pregnancy complications in humans. Because immunosuppressed organ transplant recipients are at elevated risk for developing pregnancy complications associated with altered placentation independent of allograft type, we reasoned that organ transplant recipients would provide insights into the role immune cells play in supporting placentation and pregnancy in humans. More specifically, we aimed to test the hypotheses that a specific population of innate immune cells required for normal placentation – uterine natural killer cells – was altered in organ transplant recipients, and that aberrant uNK biology in transplant recipients was due to pharmacologic immunosuppression. Since pregnancy is often discouraged in organ transplant recipients due to concerns over the well-being of the triad of mother, fetus, and allograft (*38, 39*), we leveraged access to a cohort of patients who undergo organ transplantation for the sole purpose of achieving pregnancy to address these questions. Through the comparison of multiple tissues from uterus transplant recipients with healthy controls, we identified a reduction in uterine natural killer cells in UTx patients that was associated with placental histopathology and related clinical phenotypes of placental maternal vascular malperfusion. Reduction of uterine NK cells occurred in the endometrium early after transplantation despite excellent perfusion and functional recovery of the uterine allografts, and this reduction persisted over a year in three patients whose decidua was assessed at the time of cesarean delivery. Reduction of uNKs was due primarily to loss of a subset of CD103+ NK cells which rely upon multiple NFAT-dependent molecular circuits to become tissue resident. Our work thus identifies a previously unappreciated molecular mechanism of NK cell tissue residency and links a reduction in uterine natural killer cells with altered placentation and pregnancy complications in humans.

While mechanisms of tissue residency have been well studied in infection models given the critical role that tissue resident lymphocytes play in protection against pathogens, significantly less is known about the development of tissue resident lymphocytes in other physiologic circumstances. Study of uterine natural killer cell residency programming thus facilitates our understanding of tissue residency development outside of infection, as uterine natural killer cells are required for normal placentation yet must repopulate the human endometrium after shedding during menses. Our work supports prior findings that uterine immune cells have origins in the periphery (*85*), as most immune cells recovered from the UTx biopsies possessed the recipient’s genotype. These data thus confirm not only that uterine immune cells have peripheral origins but also that immune reconstitution after transplantation is possible. However, one recipient in our cohort (UTx04) had uterine immune cells which were primarily derived from the donor in her biopsy. These findings suggest that immune reconstitution of the uterus in this patient was potentially impaired or delayed for unclear reasons. We speculate that regeneration of the superficial endometrium after the onset of menses triggers immune infiltration, and our detection of persistent donor-derived immune cells in the biopsy may signify that complete endometrial regeneration has not occurred after the stress of transplantation. Additional studies which focus on the parenchymal cell types of the endometrium will be necessary to test this hypothesis.

Our work contributes new insights to the molecular events which guide the development of tissue resident natural killer cells. Although NFAT is known to bind the *ITGAE* enhancer (*79*) and contributes to CD103 expression in CD8 T cells after TCR stimulation (*86*), a role for NFAT in regulating CD103 expression in NK cells has not yet been described. We hypothesized that in the absence of an antigen receptor, NFAT would promote CD103 expression in uterine NK cells downstream of a cytokine signal, as has been described previously for IL-6 (*82*) and IL-15 (*83*). In this work, we tested the hypothesis that IL-15 could promote CD103 expression through NFAT-dependent mechanisms given that IL-15 is secreted by decidualizing stromal cells in the secretory phase (*87, 88*). Moreover, IL-15 is known to synergize with TGF-β to induce CD103 expression on peripheral blood NK cells through MAPK pathways (*84*). Although NFAT is indeed a target of MAPK in T cells (*89*), it is unclear what intracellular molecules are promoting NFAT-dependent CD103 expression in endometrial trNK3 cells, as NFAT may be acting downstream of IL-15-mediated PI3K activation as well. Additional studies will be needed to elucidate the molecular details of NFAT signaling on CD103 expression by trNK3 cells.

Furthermore, our study suggests that human uterine NK cells are comprised of tissue resident subsets that rely on diverse mechanisms of residency programming, as we observed preferential loss of the CD103+ trNK3 subset in UTx recipients on calcineurin inhibitors that block NFAT nuclear translocation. While the loss of trNK3 cells may be a consequence of inhibition of only CD103, our data suggest additional sensitivity of trNK3 cells to NFAT blockade given their utilization of NFAT-dependent residency circuits that are upregulated in lymphocytes early after tissue entry prior to the acquisition of integrins such as CD103 or CD49a (*64, 78*). Although knock down of genes which govern early residency development in mice (e.g., *Nr4a2*, *JunB* (*78, 90*)) results in loss of CD8+ tissue resident memory cells, our work expands upon these seminal studies to demonstrate the conservation of these molecular circuits in innate lymphocyte populations in humans. Moreover, our study improves our understanding of the signaling molecules which promote early residency programming, as a role for NFAT in tissue residency has not been previously described. Whether NFAT and TGF-β synergize to promote early residency programs in addition to integrin expression remains to be determined, and it is possible that a broad range of signals in the tissue microenvironment outside of TGF-β may promote early residency. Additional studies which explore the cytokine signals leading to NFAT translocation in NK cells and other innate lymphocytes may prove critical to improve our understanding of early residency transcriptional programs. In this vein, a more important contribution of our work may be the observation that trNK1 and trNK2 subsets are preserved in the setting of NFAT inhibition and thus appear to rely on distinct NFAT-independent tissue residency circuits. Altogether, our investigations thus suggest that tissue resident lymphocytes within the same tissue can differ not only in their utilization of integrins but other types of residency programs as well. This knowledge will not only improve our understanding of disease which results from absent tissue resident lymphocytes but also help hone our ability to tune tissue resident responses to pathogens and tumors with therapeutics in the future.

Reduction of trNK subsets in uterus transplant recipients in our study cohort was associated with placental pathology and clinical syndromes associated with aberrant spiral artery remodeling. Taken together, these findings suggest that there are potentially significant consequences to disruption of tissue residency programming of NK cells on pregnancy outcomes in humans. Nevertheless, robust conclusions based on these associations are limited by several factors, including the small sample size of our UTx cohort and our inability to detect abnormalities of spiral artery remodeling (SAR) or other decidual vasculopathy. Although we did not find clear evidence of alterations of SAR in three UTx deciduas, decidual vasculopathy is a patchy process and our ability to detect decidual vascular lesions may have been impacted by our limited histologic surveillance of the decidua and the use of methodologies which do not assess the presence of trophoblast or smooth muscle cells within the vascular walls (*8*). Additional data will be needed in this regard with more comprehensive evaluation of human placentas from solid organ transplant recipients as well as in animal studies where cause-and-effect relationships can be better determined in animals exposed to NFAT inhibitors. Such studies will also be able to determine the potential contribution of the surgical procedure itself and/or alloimmunity to the development of pregnancy complications, as our study design could not address the contribution of these potential confounders to pregnancy outcomes in UTx. Animal studies and studies of additional human populations may also address another important limitation of our study, which was our inability to perform functional assays given the small amount of tissue collected. Finally, we anticipate that future studies in which serial evaluations of uterine arterial flow in pregnant UTx may help link immune perturbations in the non-pregnant endometrium with vascular consequences in pregnancy. However, it is important to note that such studies will be confounded by the inability to sample decidual immune populations directly during pregnancy, and uterine arterial Doppler studies to date have failed to predict pre-eclampsia (*91, 92*), thus explaining why this modality is not used routinely in the care of pregnant patients. Nevertheless, our data suggest that evaluation of the uterine parenchyma with Doppler microflow imaging can detect more nuanced changes over time. Studies using this approach may thus prove helpful in correlating changes in uterine profusion with alterations in immune subsets, histopathology, and/or clinical outcomes.

Moreover, while we attributed the NK reduction observed in this work to aberrant residency programming in the setting of NFAT inhibition, UTx recipients are on a variety of immunosuppression agents whose effects on trNK recruitment, survival, differentiation, or function have yet to be fully evaluated. For example, our recipients are on glucocorticoids as part of their immunosuppression regimen, and prednisone has been utilized to reduce uterine NK cells in patients with recurrent pregnancy loss (*93*). Moreover, we could not exclude a potential contribution of a systemic reduction of NK cells to our findings, as other studies have found a reduction of NK cells in the peripheral blood of kidney transplant recipients (*28, 29*). Additional studies will be needed to discriminate between these possibilities and determine how these factors play a role in endometrial immune homeostasis. Finally, our work does not preclude a role for uterine parenchymal dysfunction or altered decidualization in UTx recipients due to FK506 effects. Thus, expansion of our study to diverse cell types in other organ transplant recipients will be necessary to complete our understanding of pre-eclampsia pathogenesis in this setting. Additional examination of the NFAT axis in trNKs in other at-risk patient populations will be key to determine the potential benefit of therapeutic modulation of this pathway.

## MATERIALS AND METHODS

### Study Design

#### Research objective

To test the hypothesis that uterine NK cells are abnormal in a recipient population at elevated risk for pregnancy complications.

#### Research location

All aspects of the study (i.e., subject recruitment, sample collection, and sample analysis) were performed at the University of Alabama at Birmingham (UAB).

#### Research subjects

##### Healthy control (HC) participants

were screened for inclusion and exclusion criteria ahead of their gynecology clinic visits at UAB. Once eligibility was confirmed, participants were approached for written informed consent for study participation. Inclusion criteria consisted of reproductive age, uncomplicated pregnancy history, normal menstrual cycles, and absence of serious medical illness. Exclusion criteria included any history of the following: current pregnancy, malignancy, diabetes, pharmacologic immune modulation, systemic autoimmune disease, prior organ or bone marrow transplant, history of chemotherapy or radiation treatment, chronic or end-stage organ disease, history of HIV/HBV/HCV infection, instrumentation of the uterine cavity (i.e., D&C) within prior 12 months, use of an intrauterine device in prior 3 months, oral contraceptive use in prior 3 months, pregnancy within prior 12 months, NSAID or aspirin use within the prior 10 days, and/or medical contraindication to NSAIDs. CMV serostatus was not determined for healthy control participants. Healthy control participants underwent HCG testing in the clinical space prior to endometrial biopsy to document a negative pregnancy test.

##### Uterus transplantation

is offered as standard of care treatment outside of a clinical trial at UAB for XX genotype individuals with absolute uterine factor infertility who meet selection criteria. Uterus transplant recipients were screened for participation in the endometrial immunobiology study during clinic visits at UAB. Once eligibility was confirmed, participants were approached for written informed consent for study participation. Endometrial biopsies were collected from participants >30 days post-transplant with normal menstrual cycles and no evidence of infection. All participants were Caucasian individuals of reproductive age who met transplant center criteria; specific ages are not reported given the small nature of the patient cohort. Etiology of uterine factor infertility included hysterectomy as well as congenital absence of the uterus. PBMCs were collected one day prior to transplantation for whole exome sequencing. All participating transplant recipients received a blood group compatible transplant from a deceased donor of reproductive age. Transplant recipients received induction immunosuppression with anti-thymocyte globulin (6mg/kg) and 500mg of intravenous methylprednisolone at the time of transplantation. Maintenance immunosuppression consisted of tacrolimus, azathioprine, and prednisone. Target levels of tacrolimus were 5-10 ng/mL and were tested weekly. Secretory phase endometrial biopsies were obtained while protocol cervical biopsies were performed for immunologic monitoring of the graft. Additional medications taken during the first six months of transplantation included daily sulfamethoxazole/trimethoprim (for PCP prophylaxis) as well as acyclovir or valganciclovir for viral prophylaxis. Acyclovir was taken for HSV prophylaxis by UTx1 given that both the recipient and the donor were seronegative for cytomegalovirus (CMV). All other UTx recipients were prescribed valganciclovir as they and their donors were CMV seropositive. All endometrial biopsies from UTx recipients were obtained >30 days after transplantation after resumption of normal menstrual cycles but prior to embryo transfer (table S1). Placentas were examined immediately after cesarean delivery.

Slides from two healthy control archived placentas that were matched for gestational age (∼34 weeks and 36 weeks) with UTx deliveries were retrieved from an internal biorepository and evaluated by a perinatal pathologist (V.E.D.).

##### Brain-dead deceased donors

comprised two cohorts. Donors whose uterus was donated for the purposes of transplantation provided 1) pre-transplant blood for whole exome sequencing and 2) endometrium for genotyping controls (“Bx00”). Donors whose uterus was donated for the purpose of research provided endometrium for in vitro co-culture experiments (see table S1). Deceased donors were identified by Legacy of Hope personnel for the purpose of organ transplantation and/or research, and families of the deceased donors provided authorization. PBMCs were collected from deceased donors (n=5) prior to uterus procurement for the purposes of transplantation to perform donor genotyping through whole exome sequencing. Endometrium for in vitro co-culture experiments was obtained from deceased donor hysterectomies (n=4) that were performed during surgical training procurements but were not transplanted. Data on the menstrual cycle phase for any of the deceased donor endometrial samples was unavailable.

#### Experimental Design

##### Design Synopsis

Human endometrium was sampled from 1) healthy control participants, 2) uterus transplant recipients, and 3) brain-dead deceased donors for this study. Endometrial dating was known for all healthy control volunteers and uterus transplant recipients but not for any deceased donor sample, as healthy control individuals and UTx recipients used ovulation predictor kits as discussed below. Endometrial samples were analyzed with single cell approaches (i.e., scRNA-seq, CITE-seq, flow cytometry) as well as imaging (i.e., immunofluorescence microscopy). Single cell suspensions of endometrium were cultured with IL-15 and FK506 for 3-7 days.

##### Rigor & reproducibility considerations

Healthy control and uterus transplant participants used an ovulation predictor kit (Clearblue^®^ digital ovulation tests (SPD Swiss Precision Diagnostics GmbH; Item number 245-03-0301)) (OPK) to objectively identify the day of ovulation and thus the start of the secretory phase of the menstrual cycle. Menstrual cycle dating was therefore not dependent on subject reporting based on the start date of menses, and even individuals who were biopsied in the proliferative phase used an ovulation predictor kit during at least one menstrual cycle preceding biopsy to inform the timing of biopsy. The digital ovulation predictor kit turns positive when luteinizing hormone (LH) is detected in the urine. Participants biopsied in the secretory phase were brought into the clinic to undergo endometrial biopsy in the days following the LH surge, and biopsy data are thus reported as “LH+7” to denote secretory phase cycle day after the LH surge. Participants biopsied in the proliferative phase used the OPK to more accurately define menstrual cycle length, and then were biopsied in the cycle after detection of the LH surge. Participants still used an OPK in the cycle in which they were biopsied to ensure that biopsy had occurred prior to the LH surge. Variability in the uterus transplant data were minimized by collecting biopsies >30 days after transplantation once individuals had resumed normal menstrual cycles.

Use of Biologic and Technical Replicates: We evaluated a total of 58 endometrial, decidual, and placental samples in this work. 53 samples were collected from our laboratory, while data from five term decidual samples were downloaded for aggregation and analysis (*58*). For single-cell RNA-seq experiments, six secretory phase endometrial biopsies were obtained from six independent individuals (for healthy controls), and nine endometrial biopsies were obtained from five independent individuals who had received a uterus transplant (table S1). Five endometrial biopsies were performed on five individual deceased donors prior to procurement and transplantation. All nine UTx endometrial biopsies were taken as independent samples at distinct time points after transplant. All endometrial biopsies therefore represent biologic replicates, including two individuals who had multiple biopsies in time after transplant (UTx1 [n=4 sequential biopsies], UTx3 [n=2 sequential biopsies]). Five libraries were also prepared from three term decidual samples from three individual UTx recipients (UTx1, UTx2, UTx3), two of whom thus had technical replicates (i.e., two libraries each from UTx1 and UTx3). Results of decidual replicates were pooled for downstream analysis and reporting. Healthy control term decidual data (n=5) from Sureshchandra et al. (*58*) were downloaded and analyzed separately as well as after Harmony aggregation with decidual libraries prepared from UTx recipients as above. Endometrial biopsies from HC and UTx were similarly analyzed separately and then analyzed together after Harmony integration to control batch effects. For ex vivo flow cytometric analyses, 17 secretory phase endometrial biopsies and 8 proliferative phase biopsies were obtained from a total of 25 healthy control volunteers. For in vitro co-culture experiments, four secretory phase endometrial biopsies were obtained from healthy control volunteers and four endometrial biopsies were obtained from deceased donor hysterectomies (unknown menstrual cycle date). In vitro co-culture experiments were performed in duplicate or triplicate. For immunofluorescence microscopy analyses, secretory phase biopsies were obtained from two individual healthy control volunteers, and three secretory phase biopsies were obtained from two individual uterus transplant recipients. Although sample number was limited in this study, the entirety of the tissue was assessed with multiple sections evaluated by an automated approach with ∼10% error. Data from all individual biologic replicates are provided in the visualizations.

#### Regulatory considerations

The University of Alabama at Birmingham Institutional Review Board reviewed and approved sample collection from healthy control participants under project “Mechanisms of Uterine NK Cell Differentiation” (IRB-300006859) and sample collection from uterus transplant recipients under project “Endometrial Immunobiology in Uterus Transplantation” (IRB-3000007980). All samples were de-identified.

### Sample collection & processing

#### Endometrial biopsies

Secretory phase endometrial biopsies were collected with a Pipelle^®^ suction curette (Cooper Surgical; Ref. 8200) in the clinical setting and transported to the laboratory in DPBS with calcium and magnesium (DPBS^+/+^; Gibco™; Cat. 14-040-133) on ice. Multiple passes of the suction curette were performed to acquire sufficient tissue for multimodal analysis with processing described below. Endometrial biopsies were then digested to isolate cells for downstream analysis including sorting, flow cytometry, or in vitro incubation.

##### Endometrial biopsies used for immunofluorescence microscopy

were embedded in O.C.T. mounting media (Tissue-Tek; Cat. No. 4583) in a mold (Tissue-Tek; Cat. No. 4566). Molds were then placed in liquid nitrogen-cooled isopentane (Fisher Scientific; Cat. No. O3551-4). Frozen molds were stored at −80^°^C until sectioned onto glass slides for imaging.

##### Endometrial biopsies used for single cell isolation

were processed into single cell suspensions for use in downstream single cell isolation techniques, such as scRNA-seq, spectral flow cytometric analysis, and in vitro culture. The biopsies were minced in a petri dish containing 2-3 mL of 37^°^C-warmed DPBS with Calcium and Magnesium (DPBS^+/+^) on ice with a razor blade or scalpel and transferred to a 50 mL conical for digestion, also on ice. The petri dish was rinsed 4-5 times with 1 mL of 37^°^C-warmed DPBS^+/+^ which was also transferred to the 50 mL conical. A 1:100 ratio of 5mg/mL Liberase (Sigma Aldrich; SKU 5401127001) and a 1:1000 ratio of 1 mg/mL DNAse I (Sigma-Aldrich; SKU 11284932001) was added to the conical and inverted 2-3 times to mix. The conical containing the minced biopsy and digestion media was then placed in a 37^°^C shaking incubator at 240 RPMs and was inspected every 5 minutes to determine if the digestion had completed. If the digestion needed to continue, the conical was inverted 2-3 times to mix and placed back in the 37^°^C shaking incubator at 240 RPMs. Digestion typically took 10-15 minutes on average.

After completion of digestion, the digested sample was placed on ice and filtered through a 70 µm strainer (Stemcell Technologies; Cat. 27260) into a new 50 mL conical. The strainer was rinsed 5 times with 1 mL ice-cold DPBS^+/+^ before adding more ice-cold DPBS^+/+^ to the 50mL mark. The digested biopsy was then centrifuged at 300xg for 5 min at 4^°^C. After centrifugation, the supernatant was aspirated without disturbing the pellet, which was then transferred to a 15mL conical and resuspended in 2 mL ACK lysis buffer (Quality Biological; Cat. 118-156-101) to incubate on ice for 2 minutes. After ACK lysis, 13 mL ice-cold DPBS without calcium and magnesium (DPBS^-/-^; Gibco™; Cat. 14-287-080) was added to the cell suspension. The 15 mL conical was then inverted 2-3 times to mix and was then centrifuged at 300xg for 5 min at 4^°^C. The supernatant was aspirated without disturbing the cell pellet which was then resuspended in 0.5-1mL fresh sort buffer (ice-cold DPBS^-/-^ + 0.04% (w/v) BSA (Jackson ImmunoResearch; Cat. 001-000-162)) for counting. Cells were counted by creating a 1:10 or 1:100 dilution with 0.04% Trypan blue in PBS (Gibco™; Ref. 15250-061). For the 1:10 dilution, 10 µL of the well-mixed cell suspension was added to 90 µL of 0.04% Trypan blue in PBS. For a 1:100 dilution, 10 µL of the 1:10 dilution was added to 90 µL of 0.04% Trypan blue in PBS. Cell counts were performed using a disposable Neubauer improved hemocytometer (INCYTO; DHC-N01-5), following the manufacturer’s instructions.

#### Decidual processing

Decidua processed for <u>histologic analysis</u> was prepared from placental samples by rolling the membranes and placing into formalin. Samples were embedded in paraffin and stained for H&E.

Decidua processed for <u>single-cell RNA-seq experiments</u> was scraped from the placenta into large petri dishes containing 2-3 mL of 37^°^C-warmed DPBS^+/+^. Samples were then processed into single cell suspensions in a similar manner as the endometrial samples with modifications to accommodate the larger volume of tissue. Such modifications included: 1) using more 50 mL conical tubes for digestion, with consolidation of the sample into a single conical after being strained prior to ACK lysis; and 2) the volume of digestion media varied depending upon the size of the sample. The amount of 37^°^C-warmed DPBS^+/+^ added to the sample until the sample was fully covered by media. The amount of 37^°^C-warmed DPBS^+/+^ was tracked to ensure the 5mg/mL Liberase (Sigma Aldrich; SKU 5401127001) and 1 mg/mL DNAse I (Sigma-Aldrich; SKU 11284932001) added remained at a 1:100 and 1:1000 ratio, respectively. Finally, ACK lysis was performed twice if the cell pellet still contained significant red blood cells after the first lysis attempt.

#### Placental processing

Placentas were recovered from the operating room at time of delivery and placed in a sterile plastic container for transport to the clinical surgical pathology laboratory. Placentas were processed according to the Amsterdam Criteria (Amsterdam Placental Workshop Group Consensus) and evaluated by an experienced perinatal pathologist (V.E.D.). In brief, the placental disk was sequentially sectioned and systematic random samples of the chorionic plate, the mid-zone, and the basal plate were collected and placed in 10% formalin. Samples were subsequently embedded in paraffin, stained with hematoxylin and eosin, and sectioned at 5µm for histopathologic analysis.

#### PBMC processing

PBMCs were isolated to perform whole exome sequencing (WES). Blood was collected in EDTA-K2 tubes vacutainers (BD and Company; Ref. 366643) and blood collection set (MYCO Medical Supplies, Inc.; Ref. GSBCS23G-7T). PBMC isolation was performed at room temperature until red blood cell (RBC) lysis. Equal volumes of wash media (DPBS^-/-^ + 2% (v/v) FBS (Gemini Bio; Ref. 100106)), were added to the whole blood and mixed. The manufacturer’s instructions were followed to isolate PBMCs using Lymproprep^TM^ (Stemcell Technologies; Ref. 07811) and either a Sepmate™-15 or −50 (Stemcell; Ref. 85420 or 85450), depending on the blood volume. Once the PBMCs were isolated, the cells were washed with the DPBS^-/-^ and 2% FBS two times, centrifuged once at 300xg for 8 minutes and once at 120xg for 10 minutes with no break. After removing the final wash buffer, RBCs were lysed using 3 mL of room temperature ACK lysis buffer (Quality Biological; Cat. 118-156-101) for 2 minutes on ice. Ice-cold DPBS was added to the 14 mL mark and mixed to finish the lysis reaction. Cells were pelleted by centrifugation, 400xg for 5 minutes at 4°C. The buffer was removed and 0.5-1mL DPBS^-/-^ was added; the cells were suspended and counted. PBMCs were cryopreserved until thawed for DNA isolation and subsequent whole exome sequencing.

#### Deceased donor hysterectomies

Surgical hysterectomy for the purposes of research was performed in the Donor Recovery Center at the University of Alabama at Birmingham from deceased donors after family authorization for research was obtained. Deceased donor uteri were recovered after donor cross clamp and removal of other organs for the purposes of transplantation. Uteri were transported to the laboratory on ice and bivalved to expose the endometrium. A scalpel blade was used to gently scrape the endometrium into a culture dish, and the sample was washed and digested as above for flow cytometric and/or in vitro co-culture experiments.

### Flow cytometric analysis & cell sorting

Live, CD45+ immune cells were used for scRNA sequencing and were prepared for <u>FACS-sorting</u> from endometrium as follows: Cells were first stained using the 150 µL 1:600 LIVE/DEAD^TM^ Fixable Aqua stain kit (Molecular Probes™; Cat. L34957) diluted in DPBS^+/+^ on ice for 30 minutes. After the incubation, cells were centrifuged at 300xg for 5 minutes at 4^°^C, and the buffer aspirated. Cells were then washed in staining buffer (DPBS^+/+^ + 2% FBS + 0.4% (v/v) + 0.5 M EDTA (Fisher Scientific; Cat. No. AAJ15694AP)). Cells were pelleted and then resuspended in staining buffer and stained for 30 minutes on ice with 1 mg/mL AF700-mouse-anti-human CD45 (BioLegend; 368508) and PE-mouse anti-human CD235a in staining buffer. After antibody staining, cells were centrifuged at 300xg for 5 minutes at 4^°^C and the staining buffer aspirated. Cells were washed in 1 mL ice-cold sorting buffer and then centrifuged at 300xg for 5 minutes at 4^°^C.

The wash buffer was then removed, and the cells were resuspended in ice-cold sorting buffer at a concentration of 2-7 million cells/mL. Live CD45+ CD235a-cells were then sorted on a FACSAria I or II (BD Biosciences) at the UAB Flow Cytometry Core Facility in tubes with 5 mL sorting buffer in the bottom. Cells were counted again to determine cell number, concentration, and viability.

Cells prepared for <u>flow cytometric analysis</u> were aliquoted to have between 200,000 – 10,000,000 cells per tube. Cells were stained first in 150 µL LIVE/DEAD^TM^ Fixable Aqua using the staining kit described above. Cells were then resuspended in 50 mL staining buffer on ice, which included 30 mL of BD Brilliant Stain Buffer (BD Biosciences; Cat. No. 563794) and 1 mL 400 mg/mL Pacific Blue-mouse-anti-human CD45 (BioLegend; Cat. No. 304029; clone HI30), 1 mL 50 mg/mL PE/Dazzle594-mouse-anti-human CD16 (BioLegend; Cat. No. 302054; clone 3G8), 1 mL 100 mg/mL PE/Cy7-mouse-anti-human CD39 (BioLegend; Cat. No. 328212; clone A1), 1 mL 50 mg/mL APC-mouse-anti-human CD56 (BioLegend; Cat. No. 318310; clone HCD56), 0.5 mL 200 mg/mL APC/Cy7-mouse-anti-human CD3 (BioLegend; Cat. No. 317342; clone OKT3), 2 mL 200 mg/mL BV711-mouse-anti-human CD49a (BD Biosciences; Cat. No. 742361; clone SR84), and 1 mL 70 mg/mL BV421-mouse-anti-human CD103 (Biolegend; Cat. No. 350214; clone Ber-ACT8). After the incubation, cells were centrifuged at 300xg for 5 minutes at 4^°^C and the buffer aspirated. Cells were then washed in 1mL ice-cold staining buffer. The cells were centrifuged at 300xg for 5 minutes at 4^°^C and the buffer aspirated. Cells were resuspended in DPBS for flow cytometric analysis on a Cytek Northern Lights spectral cytometer.

### In vitro culture

Single-cell suspensions of unfractionated endometrium were spun down at 4°C, 300g, 5min and resuspended in RPMI 1640, 1X with L-glutamine & 25mM HEPES (Corning Inc., 10-041-CV), 10% Benchmark™ Fetal Bovine Serum (Gemini Bio, 100-106), and 50µM 2-ME. Culture media also included 10 ng/mL of IL-15 (Thermo Fisher, 200-15-10UG). Cells were plated in 24 well plates (Costar Corning Inc., Ref. 3738) to a final volume of 1mL per well consisting of 5×10^5^ – 1×10^6^ cells ± 10 ng/mL FK506 (Sigma, PHR1809-150MG reconstituted in DMSO) in duplicate or triplicate. All plated samples were cultured in a 5% CO_2_ incubator for 4-6 days. The culture media was replaced with fresh culture media supplemented with IL-15 as above ± 10ng/mL FK506 every 48 hours for the 4 day and 6 day cultures. Upon the cultures reaching their target dates, cells were recovered from the plates and subjected to spectral flow cytometry analysis as described.

### Whole exome sequencing

PBMCs used for whole exome sequencing (WES) were collected from uterus donors and recipients prior to uterus transplantation. PBMCs were cryopreserved, as described above, and then thawed for WES. Cryopreserved cells were partially thawed in a 37^°^C water bath while gently swirling. When half of the volume of the cells were thawed, 500 mL of 37^°^C-warmed thawing media, RPMI with 10% FBS, was added to the cryovials. Suspended cells were transferred to a 15 mL conical tube with 9 mL of warmed thawing media. Cells were centrifuged at 400xg for 5 minutes at 4^°^C. The thaw media was aspirated, and cells were washed again in thaw media. After centrifugation as before, thaw media was aspirated, and cells were suspended in DPBS^-/-^ and counted.

Genomic DNA was then isolated from 1-5 million isolated PBMCs using the DNeasy Blood and Tissue kit (Qiagen; Cat. No. 69504) following manufacturer instructions. WES libraries were prepared using the SureSelect Human All Exon V8 Kit (Agilent; 5191-6873) and sequenced on either a NextSeq 2000 (Illumina) or NovaSeq 6000 (Illumina).

### scRNA-seq and CITE-seq library preparation

Live CD45+ cells were used for single-cell RNA sequencing after being FACS-enriched as described above. Cells that were used in CITE-seq experiments were blocked with TruStain Fc block (BioLegend; Cat. 422301) for 30 minutes. Cells were then incubated with TotalSeq B™ Universal cocktail (BioLegend; Cat. 399904) in Cell Staining Buffer (Biolegend; Cat. 420201) and washed following the manufacturer’s protocol before being used for scRNA library preparation.

Cells isolated for scRNA/CITE-seq were suspended in ice-cold cell sorting buffer. scRNA GEMs and libraries were prepared according to the manufacturer’s instructions. 5’ libraries were prepared according to 10x Genomics CG000331 protocol, using Chromium Next GEM Single Cell 5’ Kit v2 (10x Genomics; PN-1000263). 3’ libraries were prepared using 10x Genomics CG000399 protocol, or CG000206 if a CITE-seq antibody panel was used with the appropriate 3’ GEM kit (10x Genomics; PN 1000121). Both 3’ and 5’ libraries used Dual Index Kit TT Set A (10x Genomics; PN 3000431). GEMs were created on either a Chromium Controller or Chromium X (10x Genomics). Either C1000 (Bio-rad) or SimpliAmp (Applied Biosystems) thermocyclers were used for library prep. Quality and concentrations of cDNA and final libraries were calculated from 2100 BioAnalyzer (Agilent) electrograms. Sequencing was performed on a NovaSeq 6000 (Ilumina).

### Data analysis

*Flow cytometry analysis*: All data were analyzed with FlowJo^TM^ software (v.10.8.1) (Becton, Dickinson, and Company; Ashland, OR, USA). Statistical analysis of flow data was performed with “stats” package in R (v.4.2.2) after importing into R. Functions used for statistical testing included wilcox.test() and the additional following arguments “paired=TRUE” or “alternative = “less”” or “alternative = “greater”” for one-sided testing for unpaired data.

#### scRNA-Seq data processing and analysis

Reads from single-cell RNA-seq experiments of each HC and UTx library were pre-processed and aligned against the GRCh38 human reference and quantified using Cell Ranger (v.6.1.1) from 10x Genomics. Downstream analysis of the generated Unique Molecular Identifier (UMI) count data was completed using R (v.4.2.1) and Seurat (v.4.4.0), utilizing default parameters unless otherwise specified. Prior to subsequent analyses, background noise from ambient RNA was removed using SoupX (v.1.6.2). The overall contamination fraction (rho) was parameterized using the autoEstCont function to remove > 2% background contamination in each dataset. The SoupX-corrected count matrixes were then loaded into R using the *Read10X* function from which Seurat objects were created per dataset via the *CreateSeuratObject* function. From each dataset, cells with less than 200 features, and features with less than 3 cells expressing them were filtered. Additionally, respective count matrixes were filtered based on quality control (QC) metrics and thresholds (table S3 and fig. S8). Doublets arising from the 10X sample loading procedure were detected and removed using Scrublet (v.0.2.3). The doublet rate per library was approximated using estimated rates provided by 10x Genomics. Similar thresholds were applied to downloaded datasets from Sureshchandra et al. (*58*).

Filtered count matrices were then merged into one Seurat object, normalized, and variance stabilized using regularized negative binomial regression via the *SCTransform* package provided by Seurat (*94*). We also regressed out variation introduced by mitochondrial expression to prevent any confounding signal. Next, identification of principal components was performed using (RunPCA). Leveraging identified PCs, and while considering each individual library as a batch, the normalized dataset was corrected for batch effects and integrated using Harmony (v1.2.0). The resulting top 50 Harmony-corrected dimensions were further used to construct a k-nearest neighbor (KNN) graph using *FindNeighbors.* Cell clustering was subsequently performed using *FindClusters* which employs a shared nearest neighbor (SNN) modularity optimization-based clustering algorithm. Resulting cluster information was used as input into the uniform manifold approximation and projection algorithm *RunUMAP* which further aided the visualization of cell manifolds in a low-dimensional space. We ran Seurat’s implementation of the Wilcoxon rank-sum test *FindMarkers* to identify differentially expressed genes in each cluster. The expression of cluster-specific canonical markers was used to annotate each cell cluster.

#### CITE-Seq data processing and analysis of CD45+ endometrial immune cells

Three healthy control and five uterine transplant datasets were further used for cell surface protein and transcriptomic data analyses. Pre-processing of these data was conducted as described previously. The RNA gene expression data was normalized using SCTransform (as earlier described), however, the ADT counts were centered log-ratio (CLR) normalized. Normalized ADT counts were then scaled and centered using the *ScaleData* function. Principal Component Analysis (PCA) was then conducted on both assays (RNA and ADT) using the *RunPCA* function with default parameters. We leveraged the ElbowPlot and DimHeatmaps to determine the number of principal components (PCs) to consider for further downstream analyses. Towards cell clustering, we found the k-nearest neighbors of each cell using the Seurat *FindNeighbors* function which then constructed a shared nearest neighbor graph using the top 40 and top 23 dimensions for RNA and ADT assays respectively. As the data constitute multiple modalities, we leveraged Seurat’s Version 4.4.0 to define cellular states based on these two assays. We applied the weighted-nearest neighbor (WNN) analysis (*95*), an unsupervised framework to integrate the multiple data types measured within each cell, and to obtain a unified definition of cellular state based on the multiple modalities. For each cell, a set of modality weights was learned, which reflects the relative information content for each data type in that cell. This enabled the generation of a WNN graph that denotes the most similar cells in the dataset based on a weighted combination of protein and RNA similarities. To achieve this, we used the Seurat *FindMultiModalNeighbors* function generating cell neighborhood graphs for further downstream analyses. We then performed cell clustering on the generated graphs and identified clusters using the Louvain algorithm (*FindCluster* function) at the default resolution. We utilized cluster information as input into the uniform manifold approximation and projection (*UMAP*) activated by the *RunUMAP* function which further aided the visualization of cell manifolds in a two-dimensional space, and as a weighted combination of RNA and protein data.

To identify differentially expressed (DE) genes between groups of cells, we used the *FindAllMarkers* function. Results from differential gene expression analysis were used to annotate clusters based on the expression of top marker genes in the resulting clusters, and by leveraging feature plots (*FeaturePlot* function) of canonical cell type specific marker expression.

#### Module scores

Module scores were calculated using the *AddModuleScore* function from Seurat. Respective feature plots were generated using *FeaturePlot* with the features set as the calculated module scores from each gene program.

#### Identification of donor or recipient cellular origin in uterus transplant samples

To determine the identity of cells from uterus transplant recipients, we used sample-specific genetic mutations and Demuxlet (v.1.0-5) (*96*). Whole-exome sequencing (WES) was conducted on DNA isolated from donor and recipient PBMC samples collected prior to transplantation. Sequenced reads were aligned using STAR. Single nucleotide variants (SNVs) were detected using Freebayes (v.1.1.0-goolf-1.7.20_39e5e4b) for each donor and recipient sample. We provided a unique identifier per sample using “bcftools reheader” and filtered the resulting VCF files to retain only common SNPs with minor allele frequency (MAF) >= 0.05, and SNPs that overlap exons (defined by GENCODE release 19). The donor and recipient VCF files were then merged and sorted to match the recipient .BAM file from cellranger using popscle_helper_tools. Reads were piled up using the popscle dsc-pileup command using default parameters along with the filtered set of barcodes from cell ranger. The genetic deconvolution of sample identities in the UTx single-cell libraries was then conducted using Demuxlet (v.1.0-5) from the popscle package by taking the merged donor and recipient .vcf file and the recipient .BAM file from Cell Ranger as inputs. Demuxlet was run at default parameters with the GT filed used to extract the genotype, likelihood, and posterior, and the offset of the genotype error rate (geno-error-offset) set to 0.05. Each cell in the final integrated dataset was annotated as donor or recipient using the BEST.GUESS column from the Demuxlet output data. Genotype results were confirmed by applying the workflow mentioned above to UTx “Bx00” samples collected from the deceased donor prior to procurement (table S1 and fig. S2) where only the donor genotype should be represented. Of note, we excluded UTx02 samples from our analysis of cell origin (Fig. 2, C and D) given concerns of cross contamination of the WES libraries based on the results of UTx02Bx00 analysis.

#### Gene set enrichment analysis

Gene set enrichment analysis was performed using the FGSEA package. Inputs included gene sets of interest and ranked differentially expressed gene lists created with MAST (*97*) and the Seurat *Findmarkers* function. Resulting differentially expressed genes were ranked based on the log-fold change (or a combinatory metric of absolute log-fold change * log transformed p-values) and the ranked gene list used as input for the fgsea R package (fast preranked GSEA), with a minimum set size of 15 genes and a maximum of 500 genes. We used the plotEnrichment() function to visualize the resulting gene set enrichment.

#### Manual curation of NFAT-dependent genes

A set of NFAT dependent genes was manually curated through literature review as well as analysis of publicly available data from NFAT-deficient mice.

### Placental Histopathology

H&E-stained slides of the placental disk were evaluated by an experienced perinatal pathologist (V.E.D.). Placental lesions were identified according to the criteria established by the Amsterdam Placental Workshop Group Consensus (*98*) and tissue was processed according to standard laboratory protocols. Placental density was calculated using automated image analysis as previously described (*99*).

### Immunofluorescence microscopy

Fresh frozen biopsies in OCT compound (Tissue-Tek) were sectioned on a cryostat at 20 mm thickness and placed on Superfrost Plus slides (Fisher) for processing. Slides were placed in 4% formaldehyde in phosphate buffered saline (PBS), pre-chilled to 4°C, for 30 minutes, then washed in PBS plus 0.3% Tween 20 three times for five minutes per wash at room temperature (RT). Slides were then dried for 15 min at RT, and sections were then treated with blocking buffer (PBS plus 1% Triton X-100, 1% normal goat serum, 1% bovine serum albumin) at RT for 1hr. Primary antibodies (rabbit anti-human NCAM1 (CD56), Cell Signaling Tech. 99746T; hamster anti-human CD3e, BioLegend 100302) were diluted (CD56: 1:50; CD3: 1:200) in blocking buffer and applied overnight at 4°C. Sections were washed in PBST three times (five minutes per wash), then treated with fluorophore-conjugated secondary antibody (donkey anti-rabbit IgG Alexa Fluor 488, Invitrogen A32790TR; goat anti-hamster IgG Alexa Fluor 594, BioLegend B388503) diluted 1:1000 in blocking buffer for 1hr at RT. Sections were then washed three times in PBST, five min per wash, then treated with DAPI (Invitrogen 62248) diluted 1:1000 in PBST for 1.5min at RT, washed once in PBST, mounted in Prolong Diamond under coverglass No. 1.5 and allowed to cure overnight. Images were obtained on ECHO Revolution in upright mode using a 20x PLAN X Apo 0.8 NA objective and a 5MP 12bit monochrome CMOS camera. Each imaged field consists of a 2416×2016 pixel array with pixel physical dimensions of 0.1777×0.1777 mm and each field covering 0.15 mm^2^ of tissue.

#### Image analysis and error estimation

Raw tagged image files were processed with custom software written in MATLAB. The analysis consisted of identifying centroids representing the centers of nuclei from the DAPI channel, then assigning cells of positive or negative CD56 or CD3 expression to each centroid. This assignment was based on relative signal in the CD56 or CD3 channel in pixels near each centroid within DAPI-based masks created as described here.

Nucleus centroid positions were estimated by a combination of iterative thresholding, difference-of-Gaussian (DoG) filtering, and iterative erosion. Single image fields were initially filtered with a Gaussian of radius 1.25 pixels, then corrected for local background changes by top-hat filtering with a disk-shaped structuring element with a 50 pixel radius. The resulting image was filtered with a disk-shaped structuring element of 0.9 pixel radius. On the resulting image, we performed iterative thresholding to find candidate "connected component" objects potentially representing single isolated nuclei or multiple overlaying nuclei. Objects whose area spanned the range of nucleus sizes determined empirically (covering between 500 and 6000 pixels) and with a circularity (calculated as four times the Area times pi divided by the perimeter) greater than 0.85 were retained as representing single nuclei. Noncircular objects were subject to iterative erosion with either a square structuring element or a diamond structuring element until either the object was divided into two or more objects, in which case two or more new centroids were taken as representing single nuclei, or the object disappeared entirely, in which case the original object was kept as a nucleus. Objects larger than 6000 pixels were subsequently processed by isolating each object’s bounding box from the filtered image and replacing all pixels outside the object with zeros. This object-specific snippet was then processed by DoG filtering with Gaussians of 10 and 12 radii. The filtered image was binarized, morphologically closed, and subjected to the same size and circularity selection criteria already described. This process was performed iteratively to subdivide all large objects into candidate nucleus objects with the selected size specificities. Centroids were then generated from all the candidate objects discovered by this method. A watershed was then applied to the original connected component mask to assign pixels to the nearest centroid. To call each centroid as expressing CD56 or CD3, the mean intensity in the CD56 or CD3 channel was assessed over all pixels assigned to each centroid after filtering with a Gaussian of radius 15 pixels. Threshold selection was performed by finding the inflection point in the rate of increase in the fraction of centroids called positive as a function of candidate threshold level.

For error estimation, we compared the computer performance to that of imaging specialists in a set of ten test images each covering 12,500 mm^2^ and containing an average of 130±33 (mean ± s.d.) hand-counted nuclei, for a total of 1,295 nuclei spanning a range of densities. Using hand-selected nuclei as the gold standard, we noted four types of error, two related to nuclear segmentation and two for assigning CD56 expression status: 1) failure to detect a nucleus, usually as a result of incorrectly merging multiple nuclei into a single nucleus; 2) counting excess nuclei, usually as a result of merging of multiple nuclei into a single nucleus; 3) incorrect splitting of single nuclei; 4) failure to call a CD56+ nucleus; 5) incorrectly calling a negative nucleus as positive. To calculate (1) and (2), we generated unique nearest-neighbor pairs of hand-selected and computer-generated centroids while using an exclusion rule prohibiting pairing of centroids separated by more than the average diameter of well-separated nuclei (8 mm). We thereby generated lists of unpaired hand-selected and unpaired computer-generated centroids and calculated the fraction of the total. Errors (3) and (4) were calculated by assessing the fraction of positive (or negative) nuclei that the computer called as negative (or positive) out of the successfully paired set of nuclei. The estimated rate for each error is 0.053±0.029, 0.051±0.043, 0.026±0.013, 0.024±0.015 (mean ± SEM). Assuming these errors arise independently, the total uncertainty in the fraction of CD56+ nuclei may be calculated by finding the root of the sum of the squares of the errors. The estimated uncertainty is thus about 8% and may be as high as 11% (0.082±0.030). Using this upper estimate, the mean fraction of cells that are CD56+ in the healthy control biopsies are 0.0303±0.0033 and 0.0357±0.0039, whereas the transplant biopsies have fractions of 0.0181±0.002, 0.0286±0.0031, 0.0125±0.0014, and 0.0122±0.0013.

### Radiographic Studies

Diagnostic <u>MRI</u> was performed per routine clinical protocols. Images were acquired pre and post administration of gadolinium contrast agent (ProHance) as indicated. <u>Pelvic ultrasound</u> images were obtained using a Philips EPIQ ELITE diagnostic ultrasound system. Transducers used were primarily a C5-1 abdominal probe for transabdominal imaging. Images were sent to the iMorgon View workstation for interpretation and QC and the Philips iSite PACS for storage. Ultrasound images were reviewed and interpreted by a diagnostic radiologist. Ultrasound imaging protocol was the same during all examinations and included the following: 1) grayscale images of the uterus in transverse and long axis, including bidimensional measurements, 2) endometrial thickness and 3) evaluation for peri-transplant and adnexal tissues. Color Doppler and MicroFlow (MFI) imaging was performed at the cranial, mid and caudal portions of the uterine allograft in both still images and cine clips. Color and spectral Doppler assessment was also performed of various vascular structures, including: 1) the external iliac artery and vein of the recipient, both proximal and distal to the arterial and venous anastomoses of the uterine graft; 2) the uterine artery at its origin from the donor internal iliac artery as well as its distal course near the mid-portion of the uterus and the lower uterine segments; and 3) the intraparenchymal arteries in the upper, mid and lower uterine segments.

## LIST OF SUPPLEMENTARY MATERIALS

Figs. S1-S8

Tables S1-S3

## ACKNOWLEDGEMENTS

The authors would like to thank the following individuals and organizations for their following contributions to this work:

1) Legacy of Hope Organ Procurement Organization
2) Uterus transplant clinical care team at the University of Alabama at Birmingham
3) Staff at the UAB flow cytometry and single cell core facilities
4) Mr. Reid Jones for support of the UAB uterus transplant program
5) Dr. Ilhem Messaoudi and Brianna Doratt for assistance with downloading data from term decidua (controls).

## Author Contributions

R.A.: data curation, analysis, visualization, software.

B.K.: data analysis; writing – original draft; conceptualization.

M.E.G.: investigation; data analysis, conceptualization.

E.W., M.B., D.E., S.D.Y., R.B., C.B., N.A., V.E.D., H.E.R., D.G.: investigation, methodology.

J.E.L.: resources, funding acquisition.

M.V.G. and B.B.: data analysis, conceptualization.

S.F., E.B., and S.C.: project administration. A.G.F.: conceptualization.

S.C.L.: conceptualization, supervision, data analysis, data curation, investigation, visualizations, writing – original draft, software.

P.M.P.: conceptualization, supervision, data analysis, visualizations, writing – original draft, funding acquisition.

All authors: writing – review and editing.

## Data & Code Availability

Raw FASTQ files have been deposited to NCBI GEO under accession number GSEXXXXXX. Code is available at: https://github.com/PorrettLab/Inhibition-of-NFAT-promotes-uterine-trNK-cell-loss-and-attendant-pregnancy-complications-in-humans

Additional data are available upon request from the corresponding author.

---

**Fig. S1.**
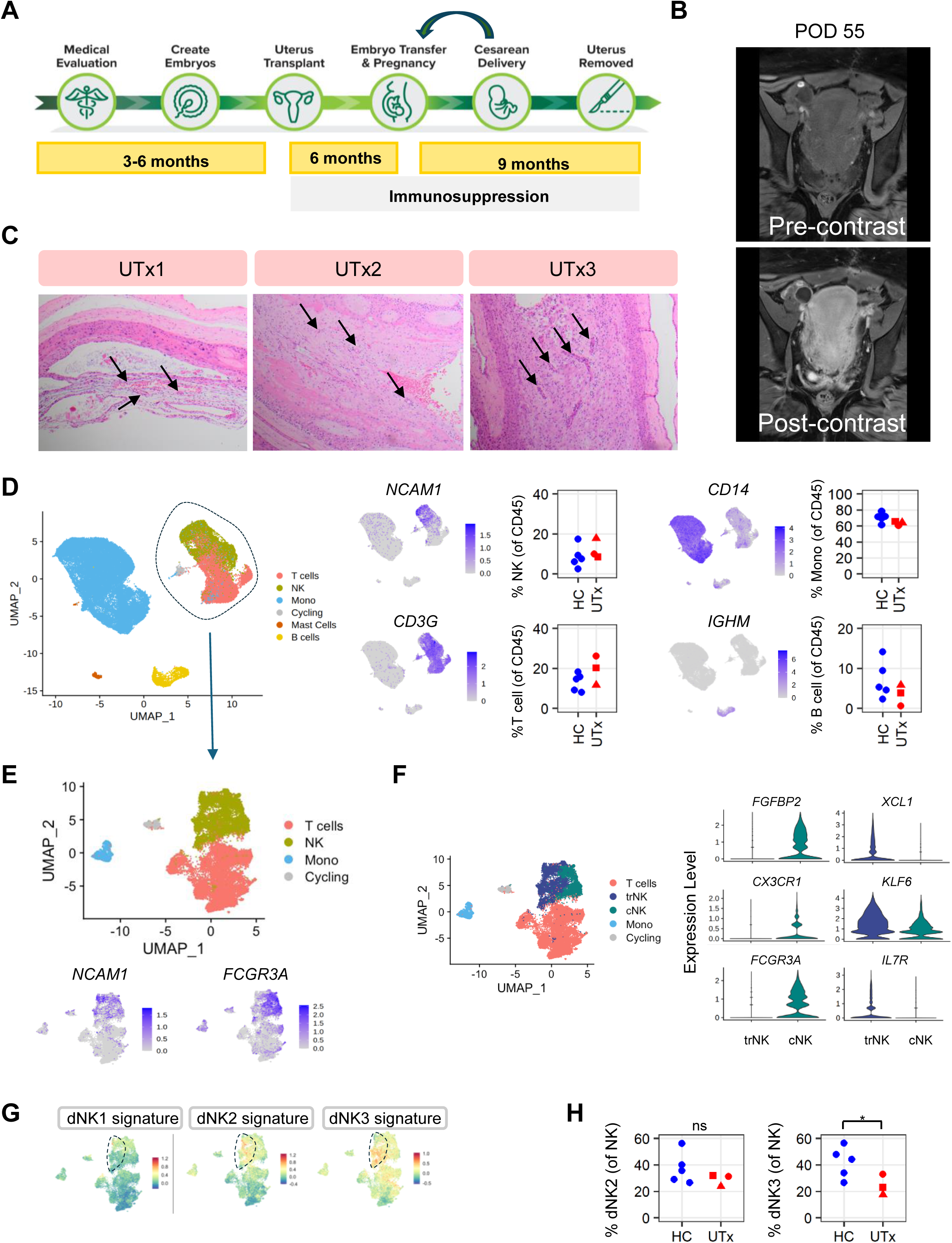
Uterus transplant overview & supplemental decidual analyses. **(A)** Summary of key events in uterus transplantation. (**B**) Images of graft perfusion 55 days after transplant (UTx2). Representative coronal T1-weighted, fat saturated MRI images of the pelvis acquired pre and post administration of IV gadolinium based contrast (ProHance) show diffuse enhancement of the allograft, confirming perfusion with no defects. (C) Normal blood vessel architecture in deciduas of UTx. Decidual vessels in the extraplacental membranes were analyzed and demonstrated normal morphology without evidence of decidual arteriopathy such as mural hypertrophy, inflammation, or acute atherosis. (H&E, 10x H&E 10X. Arrows point to cross-sections of decidual vessels. (D) *<u>Left</u>*: UMAP visualization of 64,700 immune cells aggregated from scRNA-seq of CD45+ cells sorted from three UTx term decidua (UTx1, UTx2, UTx3) (n=54,547 cells) and 5 healthy control term decidua (n=10,153 cells) (Sureshchandra et al., 2023). Dotted line encircles non-B lymphocyte cluster selected for additional analysis in E-H and Fig. 1, H and I. *<u>Right</u>*: Feature plots of marker genes discriminating major decidual immune populations and enumeration of immune subset frequency among CD45+ cells in HC and UTx. All HC vs. UTx comparisons are not significant. Circle = UTx1, Triangle = UTx2, Square=UTx3 (**E**) *<u>Top</u>*: UMAP visualization of 21,845 decidual immune cells aggregated from HC (n=5) (n=2,285 cells) and UTx (n=3) (n=19,560 cells) as in (D) and re-clustered from NK/T cluster as indicated by the arrow. *<u>Bottom</u>*: Feature plots of *NCAM1* (CD56) and *FCGR3A* (CD16) expression to delineate NK clusters in re-clustered dataset. (**F**) *<u>Left</u>*: UMAP as in (E) but re-colored to highlight cNK and trNK populations among decidual NK cells (aggregated from HC + UTx as above). *<u>Right</u>*: Violin plots of selected differentially expressed genes between decidual trNKs and cNKs. (**G**) Expression of reference first-trimester decidual NK signatures (Vento-Tormo, Nature, 2018) by term decidual NK cells aggregated and re-clustered from 5 HC and 3 UTx. Note nominal expression of dNK1 signature in term decidual NK cells. trNKs are indicated by dashed line. (**H**) Frequency of cells expressing the highest dNK2 and dNK3 signatures among all NK cells. No difference between dNK2 frequency in HC and UTx. *P-val<0.036 (one-tailed, Wilcoxon).

**Figure S2.**
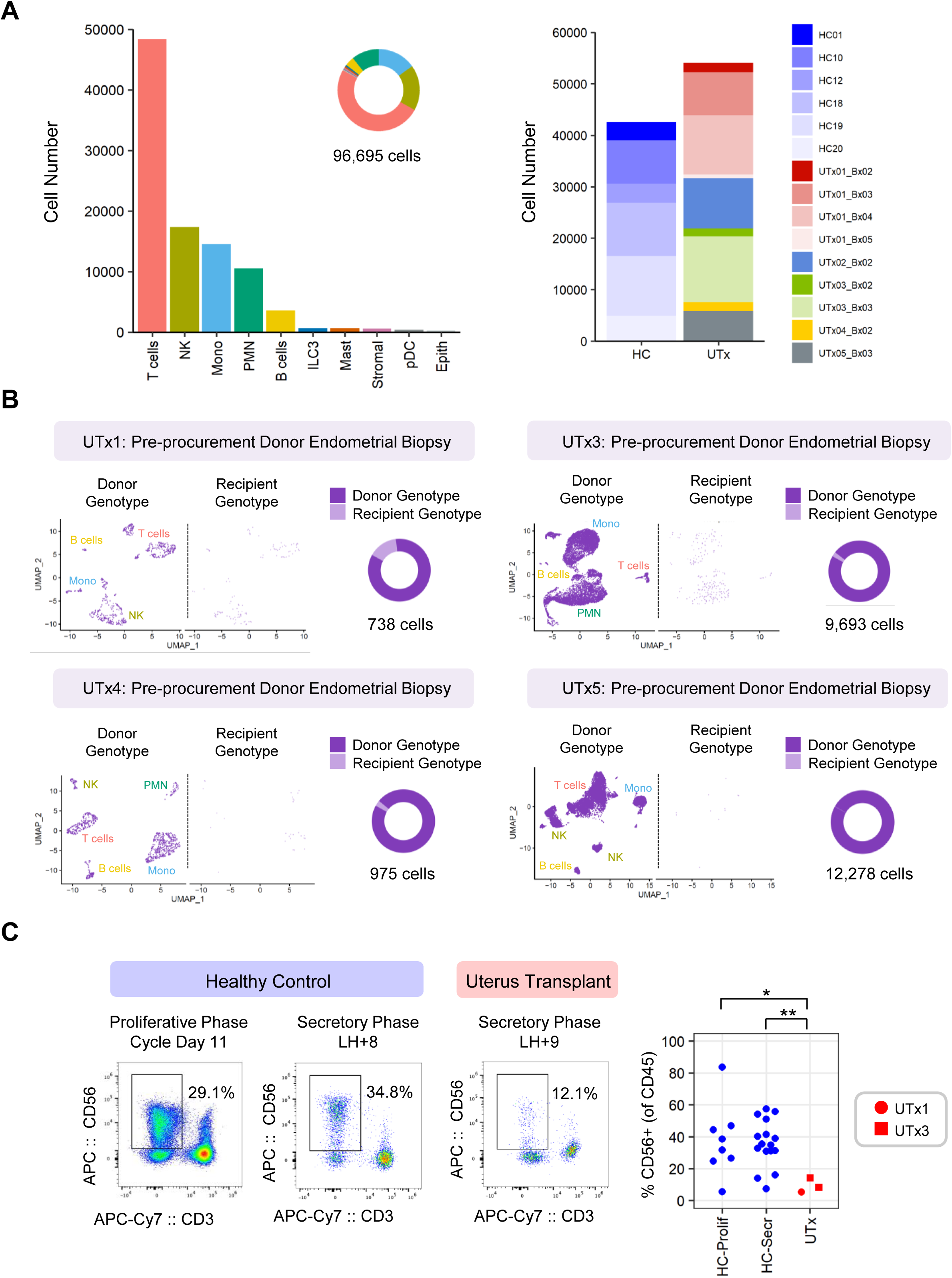
Quantification & validation of endometrial immune profiling and genotyping. (**A**) *<u>Left</u>*: Bar plot of total cell number of CD45 cells in human endometrial biopsies that underwent scRNA-seq. *<u>Right</u>*: Bar plot of total cell number by library (6 healthy controls + 5 uterus transplant recipient (9 biopsies)). Timing of UTx endometrial biopsies after transplantation is depicted in Fig. 1A and detailed in table S1. (**B**) Genotyping results of pre-procurement donor biopsies. UTx2 is not shown (see Methods). (**C**) Flow cytometry validation of scRNA-seq results in Fig. 2. Endometrial biopsies were collected from 1) healthy control volunteers in the proliferative phase of the menstrual cycle (n=8) (“HC-Prolif”; 2) healthy control volunteers in the secretory phase of the menstrual cycle (n=16) (“HC-Secr”), and 3) UTx recipients (n=2 recipients; secretory phase; n=3 biopsies with two biopsies from one individual). Biopsies were digested and single cell suspension were stained for flow cytometric analysis. Bivariate plots are gated on live, CD45+ singlets. *p<0.025; **p<0.008. One-sided Wilcoxon test.

**Fig. S3.**
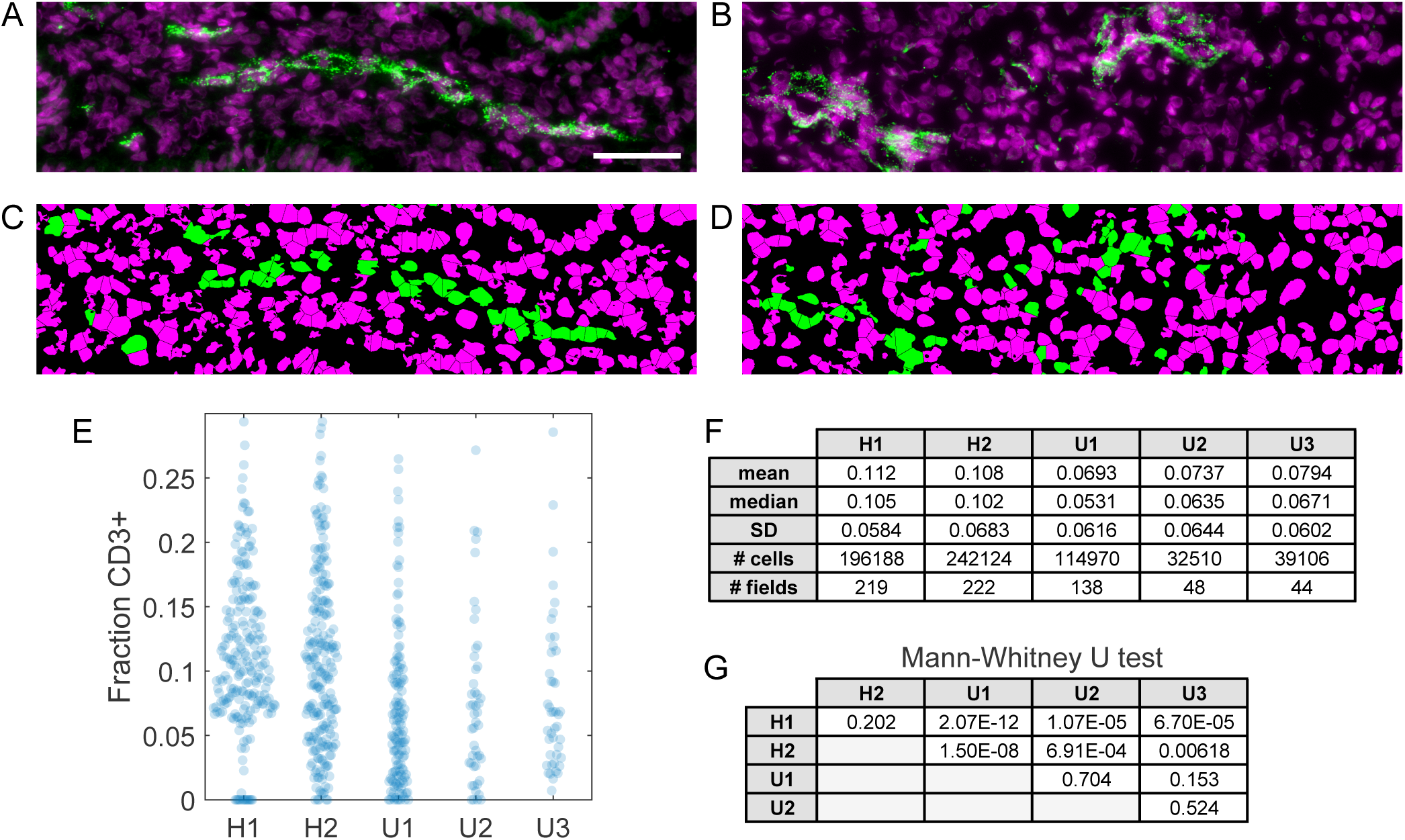
CD3-expressing cell density is reduced in transplant endometrium compared to healthy controls. (**A** and **B**) DAPI (magenta) and anti-CD3 (green) staining in healthy control H1 (A) and transplant U2 (B). Scale bar: 50µm. (**C** and **D**) Computer rendering of positive (green) and negative (magenta) nuclei in panels A and B. (**E**) Fraction of nuclei designated CD3-positive in healthy control (H1, H2) and uterus transplant (U1-U3) biopsies. (**F**) Summary statistics for the five biopsies. (**G**) p-values (Mann-Whitney U test) for all pairwise comparisons of the five biopsies.

**Figure S4.**
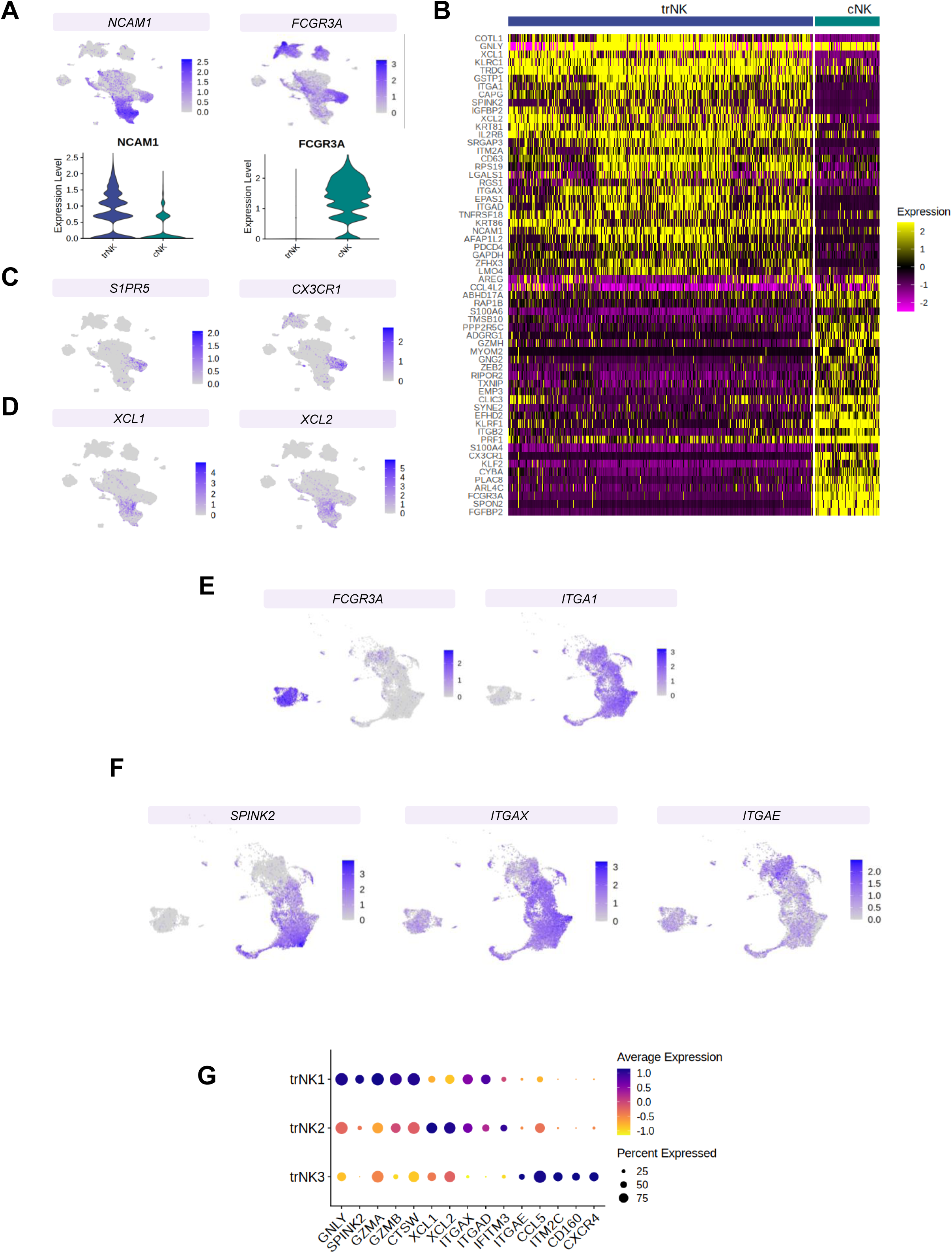
Marker genes of tissue residency or cytotoxicity and circulation in endometrial NK cells. (**A**) Feature plots of NK marker genes (*NCAM1, FCGR3A*) which correspond to CD56 and CD16 identify NK cells in the endometrium of HC and UTx. Note highest RNA expression for *NCAM1* inside the trNK population. (**B**) Differential expression analysis between trNKs and cNKs as shown. (**C**) Expression of genes relating to circulation or cytotoxic effector function in cNKs. (**D**) Expression of chemokine ligands in trNK subsets. (**E**) Reciprocal expression of *ITGA1* and *FCGR3A* in trNKs and cNKs, respectively, in NK cells re-clustered from the CD45 data. (**F**) Feature plots of key marker genes for trNK1, trNK2, and trNK3 subsets. (**G**) Bubble plot of marker genes defined for trNK1, trNK2, and trNK3 subsets.

**Figure S5.**
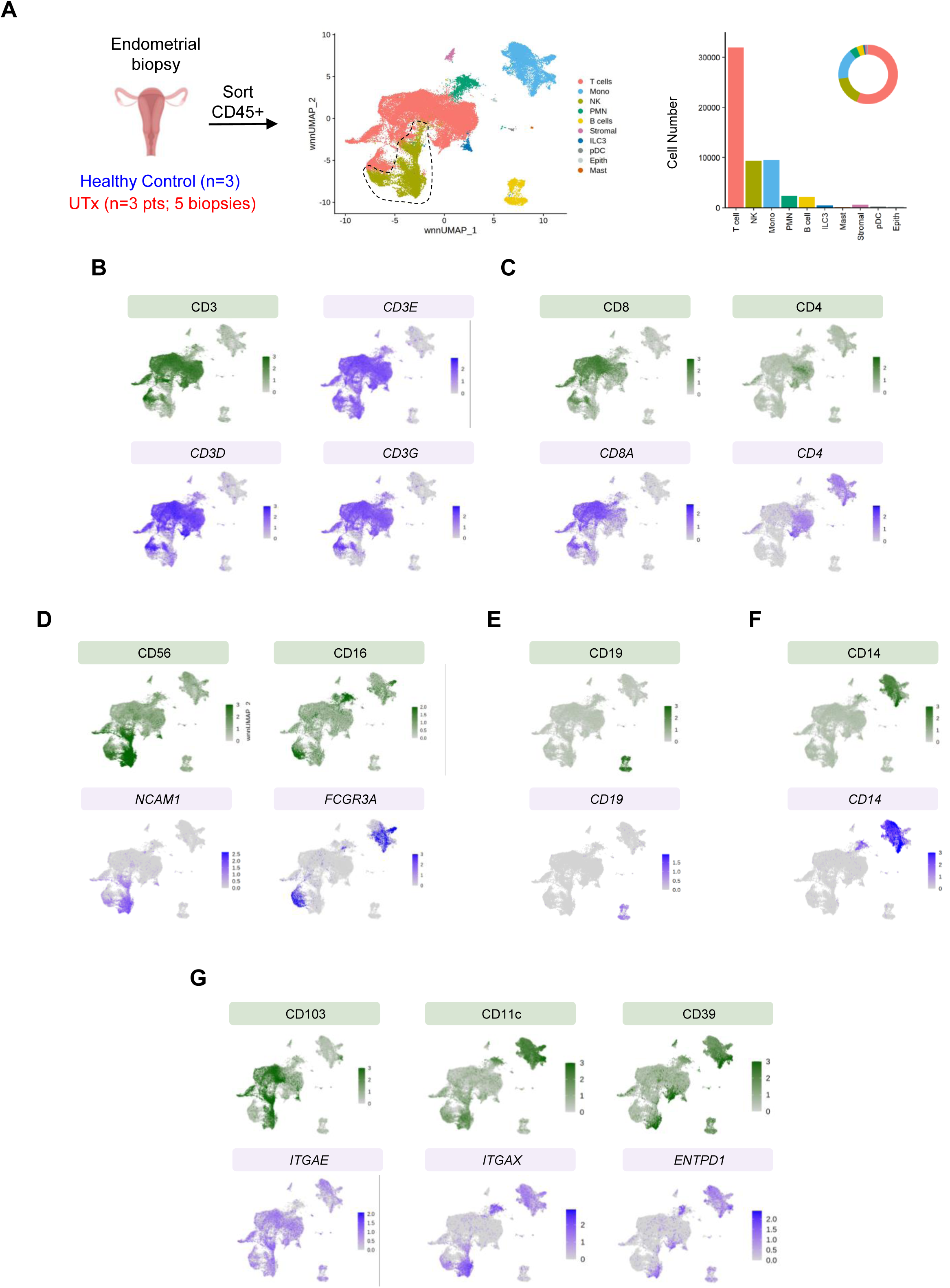
CITE-seq analysis of endometrial CD45+ immune cells from HC and UTx. (**A**) Schematic of data collection and CITE-seq analysis. Weighted nearest neighbor UMAP (wnnUMAP) visualization of 56,805 CD45+ immune cells sorted from 3 HC and 5 secretory phase endometrial biopsies from 3 individual UTx recipients. Dotted line indicates cells selected for re-clustering for Figs. 5 to 7 and figs. S6 and 7. **B-G**: Feature plots of indicated proteins (green) and genes (purple) in endometrial immune populations. T cell markers are shown in (**B** & **C);** NK markers are shown in (**D);** B cell markers are shown in **(E);** Macrophage marker is shown in **(F). (G)** shows expression of genes and proteins used to discriminate between populations of trNKs which are shared with other immune cells, including CD103 (trNK3), CD11c (trNK2) and CD39 (trNK1).

**Figure S6.**
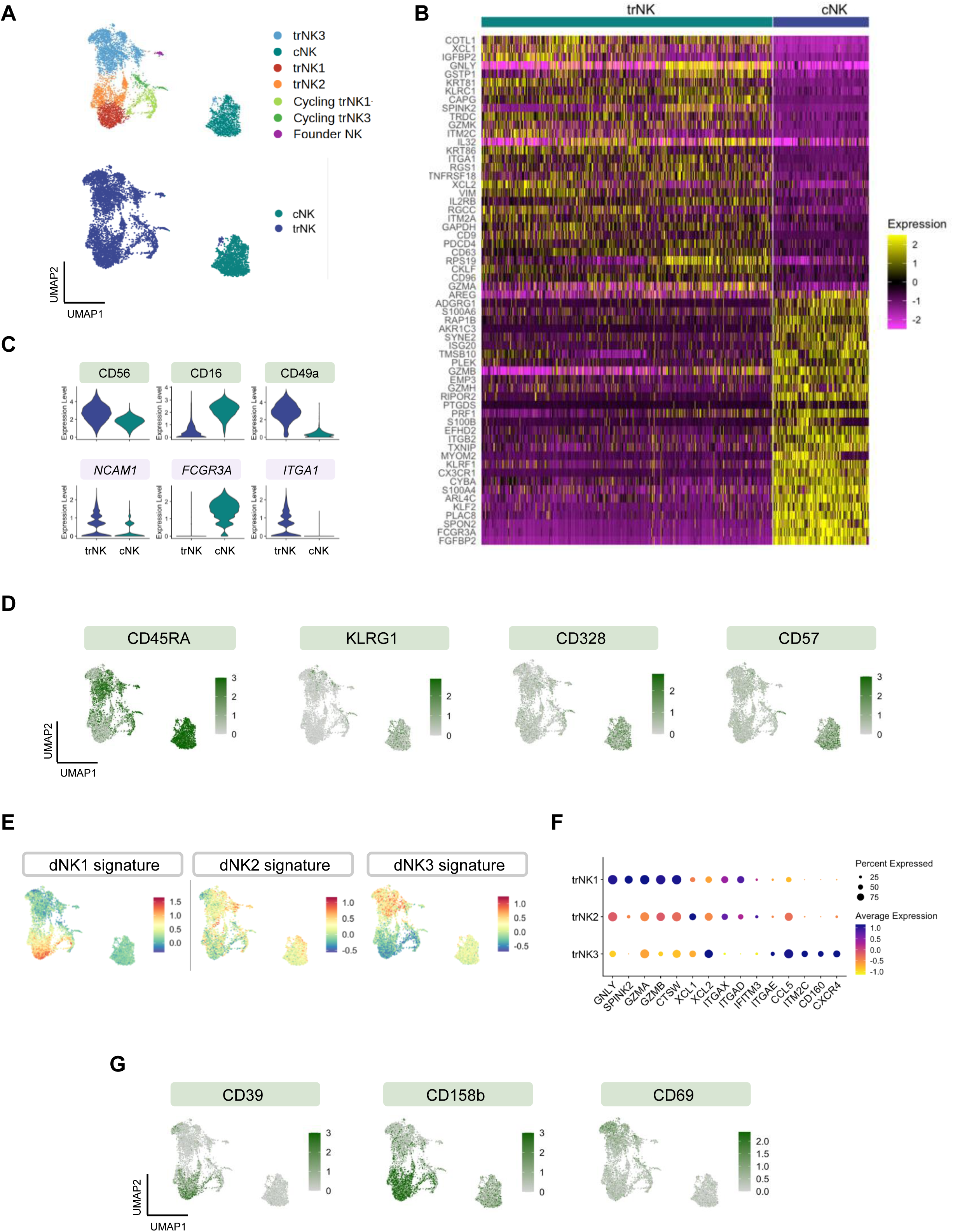
CITEseq analysis of endometrial NK cells from HC and UTx. (**A**) UMAP visualization of 6,570 NK cells re-clustered from CD45 immune cells sorted from 3 HC secretory phase endometrial biopsies and 5 UTx secretory phase endometrial biopsies (3 individual patients), after removal of NKT cells. *<u>Top</u>*: UMAP from Fig. 5 is reproduced for clarity. *<u>Bottom:</u>* NKs are categorized as trNK cells (blue) or cNK cells (turquoise) based on expression of known tissue resident genes and/or cytotoxic gene programs. (**B**) Differential expression analysis between trNKs and cNKs. (**C**) Expression of CD56 (*NCAM1*), CD16 (FCGR3A) and CD49a (*ITGA1)* on trNKs and cNKs, in NK cells re-clustered from the CD45 data. (**D**) Expression of proteins associated with cNK cells. (**E**) Module score of dNK1, dNK2, and dNK3 gene signatures on endometrial NK cells. (**F**) Bubble plot of marker genes for trNK1, trNK2, and trNK3 subsets. (**G**) Expression of proteins which distinguish trNK1s (CD39 and KIR2DL2/KIR2DL3) from trNK3 (CD69) cells.

**Figure S7.**
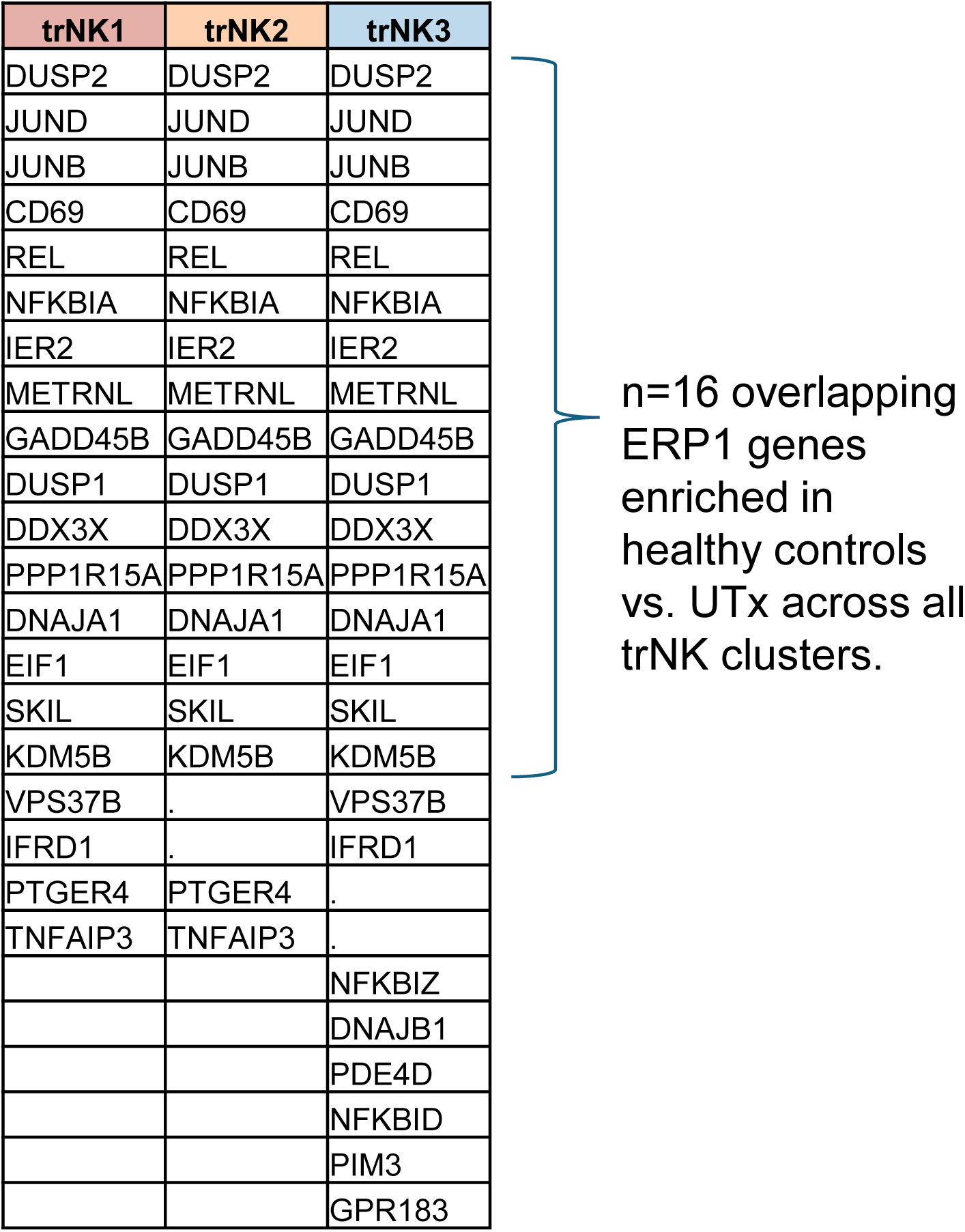
List of leading edge ERP1 genes enriched in healthy control trNK subsets (vs. UTx).

**Figure S8.**
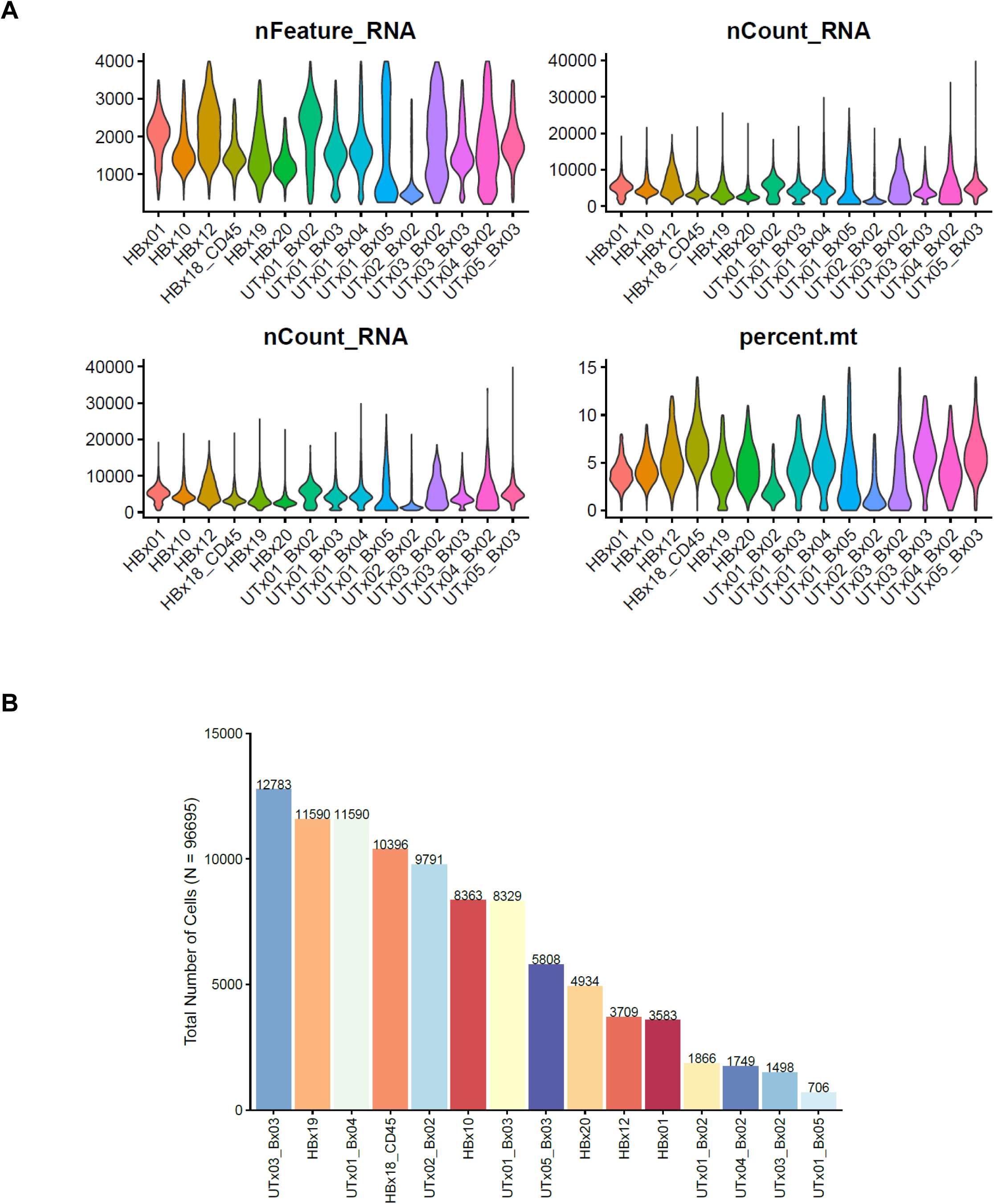
scRNA-seq QC Metrics. (**A**) Key QC metrics for scRNA-seq libraries after filtering with the thresholds as described in “Methods”. (**B**) Number of cells in each of 15 HC and UTx endometrial libraries used in the manuscript.

